# MYC-MIZ1 Complexes at Enhancers Tune Neuroendocrine Identity of Small Cell Lung Cancer

**DOI:** 10.64898/2026.08.19.745709

**Authors:** Lisa M. Fröhlich, Hannah L. Tumbrink, Bikash Adhikari, Margarita Rempe, Jenny Ostendorp, Paula Zickler, Pascal Hunold, Michaela Höhne-Wiechmann, Alena Heimsoeth, Yichun Tang, Stefanie Lennartz, Ayla Schwäbe, Lisa Werr, Matthias Fischer, Alexander Quaas, Holger Grüll, Katia Garbert, Daniela Morgenthaler, Marian Touet, Johannes A. Hildebrand, Oliver Weigert, Filippo Beleggia, Dimitrios Papadopoulos, Elmar Wolf, Johannes Brägelmann, Julia Frede, Robert Hänsel-Hertsch, Martin L. Sos

**Affiliations:** German Cancer Consortium (DKTK), German Cancer Research Center (DKFZ), Heidelberg, Germany; Department of Medicine III, LMU University Hospital, LMU Munich, Munich, Germany; Department of Translational Oncology, German Cancer Research Center (DKFZ), LMU Munich, Munich, Germany; Department of Translational Genomics, Faculty of Medicine and University Hospital Cologne, University of Cologne, Cologne, Germany; Institute of Biochemistry, University of Kiel, Kiel, Germany; Center for Molecular Medicine Cologne (CMMC), University of Cologne, Cologne, Germany; Department of Experimental Pediatric Oncology, University Children’s Hospital of Cologne, Faculty of Medicine and University Hospital of Cologne, Cologne, Germany; University of Cologne, Institute of Pathology, Faculty of Medicine and University Hospital Cologne, Cologne, Germany; University of Cologne, Department of Radiology, Faculty of Medicine and University Hospital Cologne, Cologne, Germany; Dean’s Office, Medical Faculty and University Hospital Düsseldorf, Heinrich-Heine-University Düsseldorf, Germany; Department I of Internal Medicine, Faculty of Medicine and University Hospital Cologne, University of Cologne, Cologne, Germany; Department of Hematology and Stem Cell Transplantation, University Hospital Essen, Essen, Germany; Laboratory for Experimental Leukemia and Lymphoma Research (ELLF), LMU University Hospital, Munich, Germany; Mildred Scheel School of Oncology Aachen Bonn Cologne Düsseldorf (MSSO ABCD), Faculty of Medicine and University Hospital Cologne, University of Cologne, Cologne, Germany; Mildred Scheel Nachwuchszentrum, University of Würzburg, Am Hubland, Würzburg, Germany; Theodor Boveri Institute, Department of Biochemistry and Molecular Biology, Biocenter, University of Würzburg, Am Hubland, Würzburg, Germany; University Bonn, University Hospital Bonn, Medical Clinic and Polyclinic III Internal Medicine, Oncology, Hematology, and Rheumatology; Technical University of Munich, School of Medicine and Health, Institute of Laboratory Medicine, TUM University Hospital; Cologne Excellence Cluster on Cellular Stress Responses in Aging-Associated Diseases (CECAD), University of Cologne and University Hospital Cologne, Cologne, Germany; Institute of Human Genetics, University Hospital Cologne, Cologne, Germany

**Keywords:** Small Cell Lung Cancer (SCLC), Neuroendocrine Differentiation, MIZ1, MYC, Enhancer Regulation, DNA motif binding, Chemotherapy Response

## Abstract

MYC family members have been extensively studied as undruggable transcription factors regulating oncogenic signaling in highly aggressive tumors such as small cell lung cancer (SCLC), via promoter binding. Here, leveraging the previously described Myc-driven SCLC mouse model (RPM), we generated RPM-Miz1^ΔPOZ^ (RPMM) mice to uncover a Myc-dependent regulation of neuroendocrine (NE) differentiation, via enhancers. Our functional and genomic analyses reveal that Miz1 facilitates Myc binding to low-affinity E-boxes at distal chromosomal regions, thereby enabling Myc occupancy at sites with otherwise limited intrinsic affinity. We further show that SCLC patients and cellular models share an enrichment of low-affinity E-Box Myc binding motifs at enhancer regions that loop to genes of classic neuroendocrine differentiation. Integrated epigenetic and genomic analyses with AI-modeling implicate Myc/Miz1 binding at enhancers as the determinant for the expression of bona-fide neuroendocrine genes. In RPMM tumors, the suppression of neuroendocrine identity is paralleled by a redistribution of Myc protein towards promoter-proximal regions, hyper-activation of Myc transcriptional programs, apoptotic priming and enhanced sensitivity to etoposide. Together, these findings uncover Miz1/Myc-engaged enhancers as a central hub for neuroendocrine lineage programs and provide a mechanistic basis for a targeted inhibition of Miz1 to boost chemosensitivity in SCLC.

## Introduction

Small cell lung cancer (SCLC) is an aggressive, predominantly neuroendocrine malignancy that accounts for ∼15% of all lung cancer cases and is strongly associated with tobacco exposure ^1^. A subset of SCLC seems to originate from pulmonary neuroendocrine cells (PNECs), a rare epithelial cell population in the lung involved in sensing airway oxygen levels. Recent evidence, however, indicates that additional epithelial cell types, including basal cells, can also serve as cells of origin for SCLC in specific genetic and environmental contexts^2^. Despite initial sensitivity to platinum-based chemotherapy regimens with or without immune checkpoint inhibition, most patients experience rapid relapse and develop drug-resistant disease, leading to a dismal 5-year survival rate of less than 7% ^1,3^. The absence of actionable oncogenic drivers and the high degree of intratumoral heterogeneity remain major barriers to the development of effective, targeted treatments ^4^.

On the molecular level, SCLC is characterized by ubiquitous inactivation of tumor suppressors TP53 and RB1, while amplifications of MYC family paralogs, including MYC, MYCL, and MYCN, represent recurrent oncogenic events^5–7^. These MYC-driven SCLCs are typically characterized by high proliferative signaling, a heterogenous neuroendocrine profile, and distinct therapeutic vulnerabilities ^8–10^. Transcriptional SCLC subtypes have been proposed based on the expression of transcription factors such as ASCL1 or NEUROD1 that are associated with a neuroendocrine phenotype, while POU2F3 and potentially other transcription factors may be expressed in non-neuroendocrine subtypes. However, the intrinsic heterogeneity and plasticity of cellular states limits the possibility to draw hard lines between the subtypes ^4,11,12^. MYC amplification has been specifically linked to a transition toward a low-neuroendocrine, NEUROD1-high subtype, characterized by increased cellular plasticity and reduced expression of canonical neuroendocrine markers ^13^. More recently, increased MYC occupancy at promoters of genes in deregulated pathways was observed following chemotherapy in a PDX model of SCLC ^14^. In addition, MYC has emerged as a regulator of lineage identity through enhancer-specific regulation of transcriptional programs that shape cellular fate ^15^. However, the molecular mechanisms by which MYC regulates lineage identity remain incompletely understood, particularly whether MYC functions independently or in cooperation with transcriptional cofactors. One such candidate is MYC-Interacting Zinc Finger Protein 1 (MIZ1), a transcription factor and MYC-binding partner known to have context-dependent roles in development and cancer ^16^.

MIZ1 contains a BTB/POZ (Broad-Complex, Tramtrack and Bric-à-brac / Poxvirus and Zinc finger) domain that is required for DNA binding and transcriptional repression, as well as a zinc finger domain implicated in both gene activation and chromatin binding ^17,18^. MIZ1 has classically been described as a transcriptional regulator acting at promoter-proximal regions, where it antagonizes MYC-driven gene activation through recruitment of repressive complexes ^16^. Although MIZ1 has been studied in different tumor types ^16,19,20^, its function in SCLC remains unknown. In particular, how MIZ1 influences MYC-mediated enhancer activation or repression, and whether it contributes to neuroendocrine identity or plasticity in SCLC, are key open questions.

In this study, we employed a genetically engineered mouse model (GEMM) mimicking SCLC with Myc mutation (RPM: Trp53^-/-^, Rb1^-/-^, Myc^T58A^) and combined it with a Miz1^ΔPOZ^ mutation that disrupts DNA binding while preserving non-transcriptional functions. This allowed us to uncouple Myc activity from its transcriptional co-factor Miz1 and systematically assess its impact on gene regulation, chromatin dynamics, enhancer activity, differentiation and therapeutic response.

## Material and methods

### Cell Culture

Human cell lines were obtained from ATCC and verified by STR profiling. Murine RP SCLC cell lines and spheroids were kindly provided by the laboratories of H.C. Reinhardt and F. Beleggia. Murine RPM and RPMM SCLC cell lines were derived from lung tumors of SCLC GEMMs driven by either the loss of Trp53 and Rb1 as well Myc^T58A^ point mutation (RPM) or loss of Trp53 and Rb1, Myc^T58A^ point mutation, and deletion of the BTB/POZ domain of Miz1 (Miz1^ΔPOZ^) (RPMM). RPM and RPMM lung tumors were minced and enzymatically dissociated (Tumor Dissociation Kit, mouse #130-096-730, Miltenyi Biotec) according to the manufacturer’s protocol to generate adherent cell lines. RPM and RPMM lung tumors were mechanically minced followed by enzymatic dissociation using collagenase type IV–containing tumor digestion media (100 U/mL collagenase IV, 15 mM HEPES in tumor culture media) at 37 °C for 5–30 min with intermittent monitoring to generate suspension cell lines. Adherent RPM and RPMM cell lines were cultured in cell culture flasks with standard surface, whereas suspension RPM and RPMM cell lines were cultured in cell culture flasks with suspension surface. All cell lines were routinely tested for mycoplasma contamination. PCR was performed with primers 1 and 5 ^21^ to verify the Miz1 status of the RPM and RPMM cell lines.

### *In vivo* studies

The animal studies were approved by the local animal welfare authorities. Mice were housed in a specific pathogen free barrier facility. RPMM mice were generated by crossing RPM mice (stock #029971, Jackson Laboratory) with Miz1^ΔPOZ^ mice (provided my M. Eilers, Würzburg). To induce lung tumors, mice at 8-12 weeks of age were anaesthetized with Ketavet (100 mg/kg) and Rompun (20 mg/kg) and were infected with 10^6^ - 10^8^ plaque-forming units of Ad5-CMV-Cre or Ad6-CGRP-Cre (University of IOWA) by intratracheal instillation.

Three weeks after tumor induction, mice were anaesthetized with 2.5% isoflurane and monitored weekly by MRI. A 3.0 T Philips Achieva clinical MRI (Philips Best, Netherlands) in combination with a solenoid coil designed for small animals were used for imaging. T2-weighted MR images were acquired in the axial plane using turbo-spin echo (TSE) sequence. MR images were analyzed using the Horos software. Upon detection of tumors, mice were randomized into the respective groups and treated with either vehicle or chemotherapy for a maximum of 4 cycles (5 mg/kg cisplatin once a week i.p., and 10 mg/kg etoposide 3 times a week i.p.). The experiment was ended when humane endpoints were reached or after a maximum of 28 days after treatment start.

### Cell Viability Assay

Cell viability was assessed using a luminescent cell viability assay (CellTiter-Glo, #G7570, Promega). Cells were plated in 96-well plates in technical triplicates. After 24 h, cells were treated with eight decreasing concentrations of the indicated compounds. Luminescence was measured after the specified treatment period and normalized to DMSO-treated controls. Half-maximal growth inhibitory (GI₅₀) concentrations were calculated by fitting sigmoidal dose–response curves using Prism 10 (GraphPad). Data are presented as mean ± SEM, and statistical significance was assessed using an unpaired Student’s t test.

For experiments involving inducible MYC constructs, RP1380 spheroid wildtype (WT) cells or cells stably expressing empty vector (EV), MYC-ER, or MYC^T58A^-ER were used. MYC-ER activity was induced by addition of 4-hydroxytamoxifen (OHT; 2 µM) 8–10 h prior to the start of etoposide treatment. OHT was refreshed every 24 h throughout the experiment. Cells were then treated with four decreasing concentrations of etoposide, and cell viability was measured using CTG as described above. Data are presented as mean ± SEM. Statistical significance was assessed using two-way ANOVA.

### BH3 profiling

BH3 profiling was performed as described previously^22^. In brief, cells were harvested and resuspended in MEB-P25 buffer (150 mM Mannitol, 10 mM HEPES-KOH, 150 mM KCl, 1 mM EGTA, 1mM EDTA, 0.1% BSA, 5 mM Succinate, 2.5 g/L Poloxamer 188) and incubated with BH3 peptides (BIM, BAD, PUMA, HRK-ɣ, MS1, FS1) or small molecule sensitizers (ABT-199, S63845) at final concentrations of 0.5-100 µM for 1 h. Cells were fixed with 4% paraformaldehyde for 30 mins and neutralized with N2 buffer (1.7 M Tris, 1.25 M Glycine, pH 9.1) for 20 mins. Cells were then stained with anti-cytochrome c antibody (6H2.B4) and Hoechst dye. Flow cytometry was performed using a BD Fortessa. Cytochrome c release was quantified as loss of mitochondrial cytochrome c signal, calculated based on median fluorescence intensity (MFI) and normalized to positive and negative controls. Data were analyzed using FlowJo and GraphPad Prism.

### Immunoblot

For DNA pulldown assay, biotinylated double-stranded DNA oligonucleotides corresponding to the indicated motifs (Table 1) were generated by annealing complementary oligonucleotides (95 °C for 10 min followed by incubation at RT for 1 h) and immobilized on streptavidin-coated magnetic beads (Promega, Z5481) for 1 h at RT with rotation. Beads were subsequently blocked with 3% bovine serum albumin in pulldown buffer (25 mM HEPES pH7.4, 110 mM KCl, 1 mM MgCl_2_, 0.01 mM ZnCl_2_, 10% Glycerol, 0.1% Igepal C630, 1x protease and phosphatase inhibitor cocktail (#78440, Thermofisher), 1 µM DTT) for 1 h at room temperature.

**Table 1:**
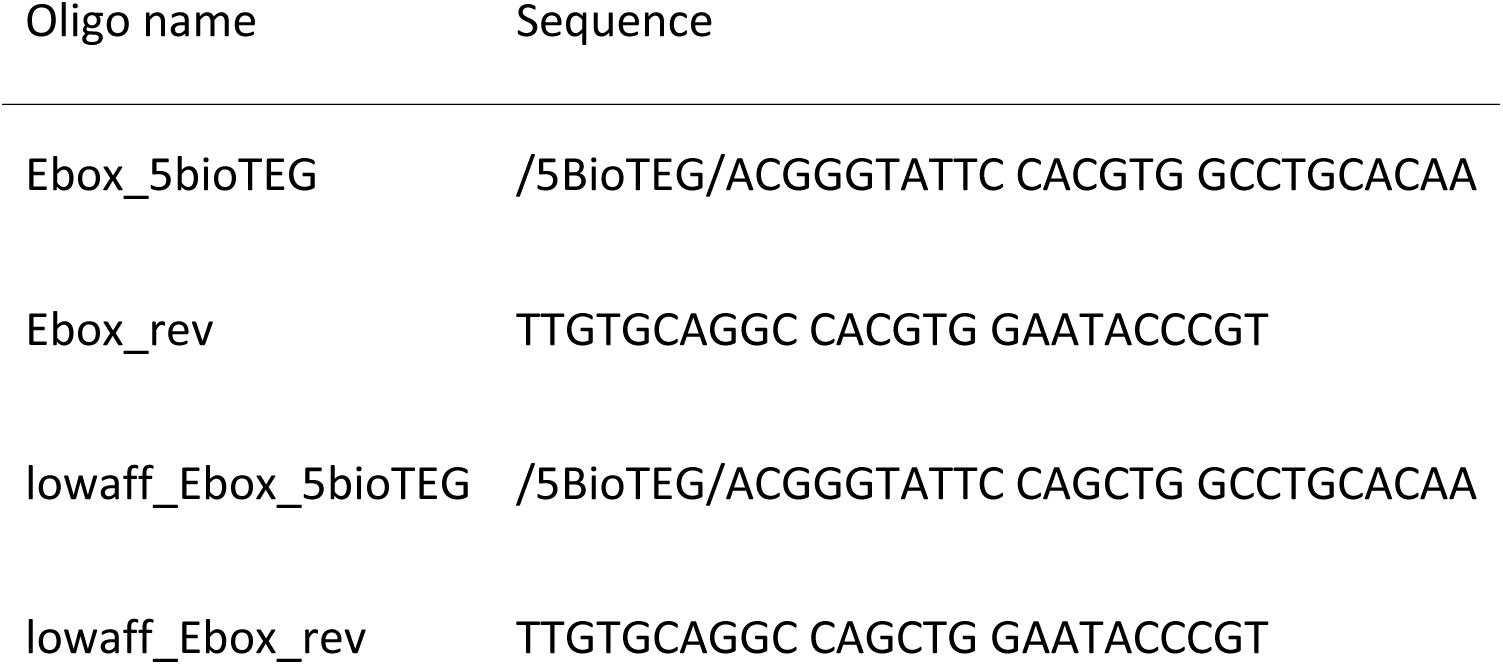
DNA-Oligos used for DNA Pulldown.

Nuclear extracts were prepared from cells using a two-step extraction protocol. Cells were harvested by scraping, washed in PBS, and resuspended in hypotonic nuclear extraction buffer (20 mM HEPES pH 7.9, 10 mM KCl, 0.5 mM Spermidine, 0.1% Triton x-100, 20% Glycerol, 1x protease and phosphatase inhibitor cocktail). After incubation on ice for 5 min, nuclei were pelleted by centrifugation (1,300 × g, 4 °C, 4 min) and washed twice in isotonic buffer (25 mM HEPES pH 7.5, 10 mM NaCl, 110 mM KCl, 1 mM MgCl_2_). Nuclei were lysed in high-salt nuclear lysis buffer (20 mM HEPES pH 7.4, 500 mM NaCl, 3 mM MgCl2, 0.2 mM EDTA, 0.5% NP40, 1µM DTT, 1x protease and phosphatase inhibitor cocktail) and incubated for 5 min at room temperature. Lysates were sonicated at 4 °C (10 cycles, 30 s on/30 s off) and centrifuged at maximum speed for 15 min at 4 °C.

For each pulldown reaction, equal amounts of nuclear protein was diluted in pulldown buffer supplemented with salmon sperm DNA and incubated with DNA-coupled beads overnight at 4 °C with rotation. Input samples (10% of nuclear extract) and negative controls lacking DNA oligonucleotides were included in parallel.

After incubation, beads were washed three times with pulldown buffer (supplemented with 300 mM KCl when high salt concentrations were indicated), and bound proteins were eluted in LDS sample buffer under reducing conditions at 70 °C for 10 min. Eluted proteins were resolved by SDS–PAGE and transferred to membranes by wet transfer. Primary and secondary antibodies were incubation for 1 h at RT. The following antibodies were used: anti-c-Myc antibody (abcam #ab32072), and as a secondary antibody Alexa Fluor 700 goat anti-rabbit IgG (Invitrogen, #A21038). Imaging was done with the Licor Odyssey DLx.

### H&E staining of FFPE tissue

Paraffin-embedded (FFPE) tissue sections were hematoxylin and eosin (H&E) stained according to standard protocols of the routine diagnostics pipeline (Institute of Pathology, University Hospital Cologne, Germany). The slides were scanned using S360 Hamamatsu Slidescanner and analyzed with QuPath.

### Immunofluorescence

For immunofluorescence experiments, RP1380 adherent cells stably expressing empty vector (EV), MYC-ER, or MYCT58A-ER were used. Cells were seeded at a density of 25,000 cells per 8-well chamber and allowed to adhere overnight. The following day, MYC-ER activity was induced by treatment with 2 µM OHT. After 6 h of induction, cells were treated with either DMSO control or 1 µM etoposide for 36 h. OHT (1 µM) was refreshed 24 h after the initial induction and maintained throughout the experiment.

Following treatment, cells were washed and fixed with 4% paraformaldehyde (PFA) in PBS for 20 min at RT. Cells were then washed three times with PBS and permeabilized with 0.5% Triton X-100 in PBS for 10 min at RT. After three additional PBS washes, cells were blocked with sterile-filtered 5% BSA in PBST for 1 h at RT.

Primary antibodies against γH2AX (Ser139 #4047974, EMD Millipore) were diluted in 1% BSA in PBST and incubated overnight at 4 °C. The following day, cells were washed three times with TBST and incubated with AlexaFluor 488 goat anti-mouse IgG (#A11001, Invitrogen) 1 h at RT in the dark. After washing three times with TBST and twice with PBS, nuclei were counterstained with DAPI (#A11001, Thermofisher) for 5 min at RT, followed by three PBS washes.

Chamber walls were removed, coverslips were mounted using mounting medium (prolong glass antifade mounting, #p36980, Invitrogen), air bubbles were eliminated, and slides were allowed to dry before imaging. Nuclear fluorescence intensity was quantified using Fiji (ImageJ).

### 3’ UTR RNA and rRNA depleted RNA sequencing

For 3’ UTR RNA sequencing (RNA-seq) of tumor tissue, fresh frozen tumor tissue was sectioned in 20 µm slices and homogenized using ceramic grinding balls. For tumor cell lines, cells were harvested and pelleted. 3’ UTR mRNA libraries were prepared using the Lexogen QuantSeq kit (#015.96, Lexogen) according to manufacturer’s protocol. Sequencing was performed on a NovaSeq sequencer (Illumina) with a 1×100bp protocol. Libraries were sequenced to a minimum depth of 12 million reads per sample.

For 3’UTR RNA-seq of FFPE tissue, each 3 biological replicates and 3 technical replicates of RPM and RPMM tumors were used. For 3’UTR RNA-seq of adherent cell lines, 6 RPM and 5 RPMM biological replicates with each 2 technical replicates were used.

For rRNA depleted RNA sequencing, RPM and RPMM cell lines were used. RNA was extracted using the Qiagen RNAeasy Mini kit (#74104, QIAGEN) according to manufacturer’s protocol. mRNA libraries were prepared using Illumina Stranded total RNA with Ribo-Zero. Sequencing was performed on a NovaSeq 6000 paired end with a 1×100bp protocol.

Raw sequencing data were aligned to the murine genome reference GRCm38 and further analyzed using the nf-core/rnaseq pipeline (v3.14.0). Differential expression analysis was performed with DESeq2 (v1.46.0) for further downstream analysis.

For quantification of enhancer RNA expression, aligned reads of the rRNA depleted RNA Sequencing was used. Read counting was performed using the Rsubread (2.20.0) package’s *featureCounts* function, with separate SAF annotation files for enhancer regions and all known coding genes (for normalization). The resulting count matrices were merged and normalized using DESeq2, and normalized counts for enhancer regions were summed per sample for downstream analysis.

### 4sU RNA sequencing

4-thiouridine (4sU)-RNA labeling and sequencing was performed as previously described ^23^. In brief, murine suspension RPM426 and RPMM575 cells were treated with 4sU (#T4509, Sigma-Aldrich) for 10 min. Two biological replicates were included. 10% human cells were spiked based on the 4sU-labeled murine cells. Total RNA was isolated using the miRNeasy kit (#217084, QIAGEN), including an on-column DNase treatment step. RNA concentration and integrity were evaluated using a NanoDrop spectrophotometer. For 4sU enrichment, equal amounts of total RNA were incubated with EZ-Link Biotin-HPDP (#21341, Pierce) dissolved in dimethylformamide (0.2 mg/ml) in biotinylation buffer (10 mM Tris pH 7.4, 1 mM EDTA) for 2 h at room temperature with continuous rotation.

Following labeling, RNA was purified by chloroform–isoamyl alcohol extraction using PhaseShield Gel Tubes (#M2302-20H, LIFESCT). The aqueous phase was recovered and precipitated by adding 1/10 volume of 5 M NaCl and an equal volume of isopropanol. Samples were centrifuged at 20,000 × g for 20 min at 4 °C. The resulting RNA pellets were washed with 75% ethanol, centrifuged again at 20,000 × g for 10 min at 4 °C, air-dried, and resuspended in nuclease-free water.

Biotinylated RNA was isolated using Dynabeads MyOne Streptavidin T1 beads (#65601, Invitrogen) by incubation in wash buffer (2 M NaCl, 10 mM Tris pH 7.5, 1 mM EDTA, 0.1% Tween-20) for 15 min at room temperature with rotation. After washing steps, labeled RNA was eluted using freshly prepared 100 mM DTT and subsequently purified with the RNeasy MinElute Kit (#74204, QIAGEN).

RNA yield was determined using the RiboGreen assay (#R11490, Invitrogen), and equal amounts of enriched 4sU-labeled RNA per sample was used for library preparation. Ribosomal RNA was removed using the NEBNext rRNA Depletion Kit (Human/Mouse/Rat; #E7400, NEB) according to the manufacturer’s protocol. Sequencing libraries were prepared using the NEBNext Ultra II Directional RNA Library Prep Kit (#E7760, NEB).

To determine the appropriate number of PCR amplification cycles, a qPCR-based approach was used. A 1:10 dilution of each library was combined with PowerUP SYBR Green Master Mix (#A25742, Thermo Fisher Scientific) and Illumina-compatible primers in 10 µl reactions, and amplification was monitored on a StepOnePlus Real-Time PCR System. The final cycle number was calculated by subtracting four cycles from the Ct value. Libraries were then amplified using 10 PCR cycles and size-selected with SPRIselect beads (#B23318, Beckman Coulter).

Next, analysis of enhancer RNA expression was performed. Reads were first separated into human- and mouse-specific reads. Size factors were calculated using DESeq2 exclusively from human spike-in reads to enable normalization across samples. SAF annotation files containing ABC-defined enhancer regions were filtered to remove duplicate genomic intervals. Read counts for individual enhancer regions were obtained using the Rsubread package (feature Counts). Differential expression analysis was then performed at the level of individual enhancer regions using DESeq2, and log₂ fold changes and adjusted p values were calculated for each enhancer region between RPM and RPMM samples. Results were visualized using volcano plots.

### Spatial Transcriptomics

Formalin-fixed paraffin-embedded (FFPE) lung tumors from three representative RPM and RPMM mouse models were processed for spatial transcriptomics. From each tumor, 2 mm tissue punches were collected and embedded into a paraffin tissue microarray mold. In total, 3 RPM and 3 RPMM punches were taken. Sections were cut from the tissue microarray and mounted onto Epredia™ SuperFrost Plus™ adhesion slides. H&E staining, probe hybridization, ligation, amplification, and library preparation were performed according to the 10x Genomics Visium HD FFPE protocol (ref A). Libraries were sequenced on an Illumina NovaSeq S1 flow cell (v1.5) using paired-end sequencing according to the manufacturer’s recommendations.

High-resolution H&E images were acquired and aligned to the Visium HD capture areas using Space Ranger (v4.0.1; 10x Genomics). Cell segmentation was performed using bin2cell (v0.3.3)^24^, integrating histological information from the H&E image. Cells were filtered based on detected gene numbers and mitochondrial transcript fractions. Cells failing basic quality thresholds were excluded prior to downstream analysis, as were cells flagged as doublets using both Robust Cell Type Decomposition (RCTD) and scDblFinder (v1.24.0). Cell type annotation was performed using RCTD with a curated mouse lung single-cell RNA-sequencing reference atlas (Azimuth LungMAP/Mouse Lung CellRef Seed). Gene expression data were normalized using *SCTransform* (v5.3.0) as implemented in Seurat ^25^.

Cell cycle state, MYC pathway activity and ABC enhancer–associated gene activity were quantified using the same approaches as described for the snRNA-seq analysis.

### Isolation of nuclei

Nuclei were either isolated from snap-frozen tumor tissue or in vitro cultured cell lines as previously described and used for single-nuclei RNA sequencing (snRNA-seq), DynaTag, and ATAC-seq experiments ^14^.

For isolation of nuclei from snap-frozen tumor tissue, samples were thawed on ice and minced with a scalpel before transfer into gentleMACS M Tubes (Miltenyi, 130-093-236) containing 3 mL L1 buffer (10 mM Tris-HCl pH 8.0, 5 mM CaCl₂, 3 mM magnesium acetate, 2 mM EDTA, 0.5 mM EGTA, 1× Complete Protease Inhibitor (#11836170001, Roche), 1 mM DTT, and 0.1 mM PMSF. Homogenization was performed using the ‘Protein-M-Tube-1.0’ program. Subsequently, 3 mL L2 buffer (L1 supplemented with 0.4% Triton X-100) were added, and the suspension was filtered through a 40 µm cell strainer. Nuclei were pelleted by centrifugation at 450 × g for 10 min at 4 °C with the brake set to ∼30%. The pellet was resuspended in 1 mL L3 buffer (L1 supplemented with 0.2% Triton X-100), and 3 mL sucrose buffer (1 M sucrose, 10 mM Tris-HCl pH 8.0, 3 mM magnesium acetate) were carefully underlaid using a 5 mL syringe. Following centrifugation at 450 × g for 10 min at 4 °C with the brake set to ∼30%, the supernatant was removed without disturbing the interface, and nuclei were resuspended in the appropriate buffer for downstream applications. For single-nuclei RNA-seq, all buffers were supplemented with 120 U/mL murine RNase inhibitor (#M0314L, NEB).

For isolation of nuclei from in vitro cultured cells lines, cells were resuspended in NE1 buffer (20mM HEPES-KOH pH7.9, 10 mM KCl, 0.5 mM Spermidine, 0.1% Triton X-100, 20% glycerol, 1x Complete Protease Inhibitor (#11836170001, Roche) and incubated on ice for 3 min. Isolated nuclei were subsequently fixed in PBS containing 0.1% formaldehyde for 1 min at room temperature with gentle agitation. Fixation was quenched by addition of 74 mM glycine, followed by centrifugation at 450 x g for 3 min at 4°C to pellet nuclei.

### Single nuclei RNA sequencing

After nuclei isolation (as described above), nuclei were counted using trypan blue staining and fixed with the Evercode Nuclei Fixation v2 Kit (Parse Bioscience, ECF2003) according to the manufacturer’s instructions. Fixed nuclei were stored at −80 °C until library preparation. Single-nucleus RNA-seq (snRNA-seq) libraries were generated using the Evercode WT Full Kit (Parse Bioscience, ECW02130) or the Evercode WT Mega Kit (Parse Biosciences, ECW02030) following the manufacturer’s protocol. Libraries were sequenced in paired-end mode on an Illumina NovaSeq 6000 platform using the NovaSeq 6000 SP Reagent Kit v1.5 (200 cycles) (Illumina, 20028401) with 5% PhiX Control v3 spike-in (Illumina, FC-110-3001). Sequencing data were processed and demultiplexed into single-nucleus expression profiles using the Split-Pipe pipeline provided by Parse Biosciences (v1.0.6).

Filtered gene-barcode matrices generated by the PARSE pipeline were imported into Seurat (v5.3.0) ^25^. Cells with >100 detected genes, >200 UMIs, <100,000 UMIs, and ≤20% mitochondrial transcripts were retained. For normalization and integration, the QC-filtered object was split by sample and each sample was processed with *SCTransform*. Integration was performed using Seurat SCT-based integration followed by shared nearest neighbor graph construction, and Leiden clustering. Cell type annotation was performed using the Azimuth Mouse Lung Cell Reference (LungMAP/Mouse Lung CellRef Seed).

Cell cycle states were inferred using canonical S-phase and G2/M-phase gene sets. Human cell-cycle gene lists were converted to mouse orthologs using biomart (2.62.1) and used for scoring. Cells were assigned to G1, S, or G2/M phase using Seurat’s CellCycleScoring function. Myc pathway activity was quantified using curated MSigDB hallmark gene sets. Per-cell Myc target activity was quantified using Seurat’s *AddModuleScore* function on SCTransform-normalized tumor cell expression data, using either mouse orthologs of HALLMARK_MYC_TARGETS_V1 or a custom ABC-derived gene set.

### DynaTag sequencing

DynaTag was performed as previously described ^14^. Nuclei were isolated according to the procedure described above and 200000 nuclei per sample were used. Next, nuclei were incubated with primary antibodies (MYC (#9402, Cell Signaling, 1:100), Miz1 (Santa Cruz, sc-136985, 1:50) H3K27ac (#MABE647, Merck, 1:100), H2A.Z (Active Motife, #39013, 1:100), H3K36me3 (Active Motife, #61101, 1:100)) overnight at 4°C, followed by incubation with secondary antibodies (0.5 µg anti-rabbit (#72763, Novus Biologicals) and 0.5 µg anti-mouse (#ab46540, Abcam)) for 1 h at room temperature, and subsequent incubation with pA-Tn5 for 1 h at room temperature. All steps were performed in AB buffer (25 mM HEPES pH 7.5, 110 mM KCl, 10 mM NaCl, 1 mM MgCl₂, 0.01% (w/v) digitonin, 1% BSA, 0.5 mM spermidine) on a nutator.

Tagmentation was performed for 1 h at 37 °C in AB buffer supplemented with 10 mM MgCl₂ using washed nuclei. For DNA extraction, 20000 nuclei were incubated in SDS- and proteinase K-containing denaturation buffer for 3 h at 58°C. Extracted DNA was further used for library preparation with custom Nextera primers. Following double-size selection cleanup, libraries were paired-end sequenced on an Illumina NextSeq 1000 platform using standard Nextera primers (IDT) and 0.5% PhiX Control v3 (Illumina, #FC-110-3001). Experiments were performed in biological triplicates with each 3 technical replicates.

Raw sequencing reads (FASTQ) were adapter-trimmed using cutadapt2 and subsequently aligned to the reference genome (mm10 or hg38) with bwa-mem2 (v2.2.1). To account for sequencing depth differences, aligned reads were downsampled to 10000000 reads to equalize library sizes across samples. Resulting BAM files were sorted and indexed, and CPM-normalized signal tracks were generated using bamCoverage. Counts were normalized using library size–based scaling, and differential peak analysis between groups was performed using a recursive comparison strategy implemented in edgeR. Merged bigWig files were generated by first merging BAM files using samtools (v1.13) merge, followed by indexing and conversion to bigWig format using bamCoverage. Peak calling was performed with SEACR (https://github.com/FredHutch/SEACR) (v1.3) using a threshold of the top 1% AUC. Reproducible peaks for each biological condition were defined by intersecting replicate-specific peak sets with bedtools multiIntersectBed, retaining only regions present in all three replicates. A unified consensus peak set was generated by concatenating reproducible peaks across conditions, followed by sorting and merging of overlapping intervals. Read counts across the consensus peak set were quantified for each sample using bedtools coverage, and the resulting values were compiled into a peak-by-sample matrix for downstream analysis.

Motif enrichment analysis was performed on genomic regions showing loss of both Miz1 and Myc binding in RPMM tumors. For HOMER-based analysis, intersected regions were resized to 200 bp and analyzed using the HOMER software (v5.1) suite with the mouse genome (mm10) as background. Known motif enrichment was assessed using default parameters, and ranked according to the p-values.

### Assay for transposase-accessible chromatin by sequencing (ATAC-seq)

ATAC-seq was carried out based on the Omni-ATAC protocol and as previously described ^14,26^. Nuclei were isolated according to the procedure described above and 200000 nuclei per sample were further used. Nuclei were washed once with ATAC-RSB (10 mM Tris-HCl pH 7.4, 3 mM MgCl₂, 10 mM NaCl in water) supplemented with 0.1% Tween-20. A total of 50,000 nuclei were used and adjusted to a final volume of 100 µL with ATAC-RSB containing 0.1% Tween-20. Following centrifugation, nuclei were resuspended in transposition mix (20 mM Tris-HCl pH 7.6, 10 mM MgCl₂, 20% dimethylformamide, 0.01% digitonin, 0.01% Tween-20) supplemented with 100 nM Tn5-i7-i5 complex (#C01070012, Diagenode, Illumina Tagment DNA Enzyme). Tagmentation was performed for 30 min at 37°C. DNA was purified using the Zymo DNA Clean and Concentrator-25 kit (Zymo, #D4006). Next, library preparation with custom Nextera primers was performed with eluted DNA, followed by a final cleanup using a 1.3× volume of SPRI beads. Libraries were paired-end sequenced on an Illumina NextSeq 1000 platform with 0.5% PhiX Control v3 (#FC-110-3001, Illumina). Experiments were performed in technical triplicates with each 3 biological replicates.

ATAC-seq raw data were processed as described above for the DynaTag experiments.

### Enhancer - Gene Interaction Prediction and Super-Enhancer Analysis

Enhancer - gene interactions were predicted using the Activity-by-Contact (ABC) model (v1.0) as previously described ^27^ and implemented using the publicly available ABC pipeline (https://abc-enhancer-gene-prediction.readthedocs.io). The ABC model scores enhancer-gene pairs based on the product of enhancer activity, measured by chromatin accessibility, and enhancer-gene contact frequency, estimated using a power-law model (gamma = 1.024, scale = 5.959, pseudocount distance = 5,000 bp), as Hi-C data were not available. ABC score thresholds were determined automatically based on the input data distribution.

For murine RPM tumor samples, ABC models were run using ATAC-seq signal only against the mm10 reference genome. Gene coordinates were derived from custom collapsed RefSeq gene bounds (UCSC mm10 refGene annotation), with TSS regions defined as ±500 bp around the annotated TSS. The top 150,000 ATAC-seq peaks, extended 250 bp from summit (500 bp total candidate region width), were used as candidate enhancer regions. Blacklist regions were defined using the ENCODE mm10 consensus signal artifact regions (wgEncodeMm10ConsensusSignalArtifactRegions.bed). Quantile normalization of enhancer activity scores was applied across samples.

For human SCLC (SCLC21H and DMS273) and NSCLC (PC9) cell lines, as well as publicly available datasets (GSE230649), candidate enhancer regions were defined using combined ATAC-seq and H3K27ac Chip-seq (for GSE230649) or DynaTag (for SCLC21H, DMS273, and PC9) signals against hg38, using the official hg38 reference files provided and otherwise identical parameters to the murine analysis. For patient-derived material (GSE281523), ABC models were run using consensus ATAC-seq peak sets derived from merged FFPE samples (n=2) and PDX samples (n=4, requiring peaks present in at least 2 of 4 samples).

ROSE-style ranking of enhancer regions was performed based on chromatin accessibility. ABC model-predicted enhancer regions were used as input and first filtered to exclude regions overlapping ±2 kb around annotated transcription start sites (TSS; mm10). Remaining regions were stitched within a maximum distance of 12.5 kb to generate composite enhancer domains. ATAC-seq signal was quantified for each stitched region and normalized to reads per kilobase. Stitched enhancer regions were then ranked by decreasing ATAC-seq signal intensity. Super-enhancers were defined based on the inflection point of the ranked signal distribution, determined using a unit invariant knee (UIK) method. Regions above this threshold were classified as super-enhancers, while all remaining regions were classified as typical enhancers.

### AlphaGenome fine-tuning and silico mutagenesis

To assess the sequence determinants of MYC and MIZ1 binding, we fine-tuned the AlphaGenome ^28^ sequence-to-signal foundation model on our own chromatin profiling data. The pretrained AlphaGenome backbone was used as a frozen feature extractor, while a custom prediction head was trained to predict two genomic signal tracks: RPM_Myc and RPMM_Myc. Input sequences consisted of fixed 16,384 bp genomic windows from the mm10 genome. Training, validation, and test sets were defined using chromosome-based splitting, with chromosomes 1–16 used for training (250,000 random regions), chromosome 17 for validation (10,000 random regions), and chromosomes 18–19 for independent testing. Signal values were obtained directly from CPM-normalized bigWig tracks generated from DynaTag experiments. Model performance was evaluated on held-out test regions by calculating mean squared error (MSE), Pearson correlation, and Spearman correlation between observed and predicted signal profiles for each track.

To evaluate the functional contribution of candidate binding motifs, in silico mutagenesis was performed using the fine-tuned AlphaGenome model. Genomic regions corresponding to Miz1-down/Myc-down loci were extracted and centered within 16,384 bp sequence windows. For analysis of MYC binding, low-affinity E-box motifs (CAGCTG) were replaced with a scrambled sequence (CTATAG). Wild-type and mutated sequences were independently scored using the fine-tuned model, and changes in predicted signal were quantified within a 400 bp window centered on the mutated motif. For each mutation, the mean change in predicted signal between reference and mutated sequences was calculated and aggregated across all motif instances.

### Analysis of publicly available datasets

Publicly available sequencing datasets were obtained from the Gene Expression Omnibus (GEO) database. Human SCLC cell line data comprising ATAC-seq, and MYC-family ChIP-seq experiments were obtained from GSE230649. RNA-seq data from human SCLC cell lines were generated previously by our group and have been published elsewhere ^9^. Murine B-and T-cell lymphoma RNA-seq data were obtained from GSE120001 ^29^. ATAC-seq data from human patient material (PDX and FFPE) was obtained from GSE281523 ^30^. ChIP-seq data were preprocessed using the nf-core/chipseq (v.2.1.0) or the nf-core/atacseq (v2.1.2) pipeline and peaks were called using MACS2. ATAC-Seq data were processed using the nf-core/atacseq (v2.1.2) pipeline and peaks were called using MACS2. For the GSE281523 dataset specifically, FFPE biopsies from two patients were merged to generate a consensus peak set. For PDX samples (n = 4), peaks were retained if present in at least two out of four samples. These consensus peak sets derived from FFPE and PDX samples were subsequently used as input for ABC model calculations. Bulk RNA-seq data were processed and analyzed using the same computational pipelines and parameters as described above for in-house generated data.

## Results

### Miz1^ΔPOZ^ Mutation Alters Tumor Characteristics of RPM Mouse Model

Both MYC and its key binding partner MIZ1 are critical regulators of tumor development (Supplementary Figure 1A) and to systematically study their interaction in SCLC we focused on the disruption of MIZ1 chromatin binding ^16^. To test whether Miz1^ΔPOZ^ could disrupt Myc signaling in SCLC, we generated a mouse model based on the RPM (Trp53^-/-^, Rb1^-/-^, Myc^T58A/T58A^) murine system that closely resembles human SCLC ^8^. Crossing RPM mice with Miz1^ΔPOZ/ΔPOZ^ mice ^21^ results in the RPMM (Trp53^-/-^, Rb1^-/-^, Myc^T58A/T58A^, Miz1^ΔPOZ/ΔPOZ^) murine system (Figure 1A, Supplementary Figure 1B). The deletion of the BTB/POZ domain in Miz1 abrogates its ability to bind DNA, thereby impairing its function as a transcription factor, while preserving its other protein-protein interactions and non-transcriptional functions ^19,31^. Kaplan-Meier survival analysis revealed a non-significant trend toward prolonged survival in RPMM mice compared to RPM controls (median survival: 66 vs. 58 days, p = 0.1158), as assessed in tumors initiated by targeted recombination in PNECs using adenoviruses carrying Cre driven by a neuroendocrine calcitonin gene-related peptide (CGRP) (Figure 1B) ^8^. Consistent with this trend, representative MRI images show a tendency toward reduced tumor burden in RPMM mice (Figure 1C). Similarly, no difference in survival between RPM and RPMM mice were observed in tumors induced by ubiquitous epithelial recombination using a CMV-driven Cre virus (median survival: 43 vs. 43 days, p = 0.91; Supplementary Figure 1C). Also, no significant difference in the number of liver metastases was observed between RPM and RPMM tumors (Supplementary Figure 1D).

**Figure 1:**
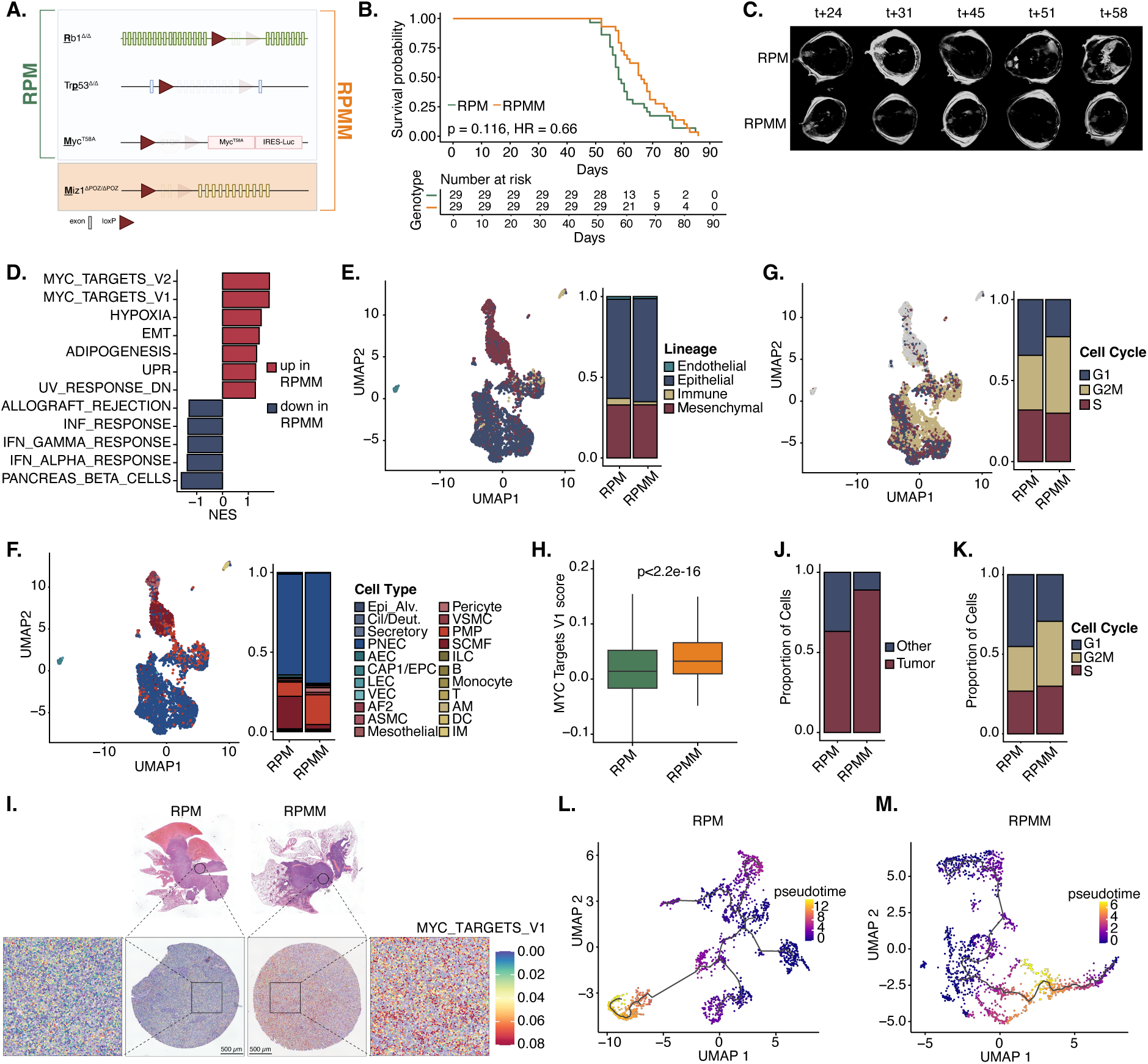
**A.** Schematic illustration of the genotype of the RPMM mouse model. RPMM harbor combined Rb1 and Trp53 deletion, MycT58A expression, and loss of the Miz1 POZ domain (Trp53^-/-^, Rb1^-/-^, Myc^T58A^, Miz1^ΔPOZ/ΔPOZ^). **B.** Kaplan Meier Analysis of mice bearing RPM and RPMM tumors. Time after infection with CGRP-Cre virus is shown. Survival differences between RPM and RPMM tumors were evaluated using the Mantel-Cox log-rank test. **C.** Representative MR images of one RPM and one RPMM mouse. The representative images were acquired at the indicated time points (t, days after tumor induction). **D.** Gene Set Enrichment Analysis (GSEA) of Hallmark pathways of 3’UTR bulk RNA-seq of FFPE tissue isolated from RPM and RPMM tumors. Pathways with p-value < 0.05 were included. **E.-G.** Relative lineage (E.), cell type (F.), and cell cycle phase (G.) composition of snRNA-seq profiles from RPM and RPMM tumors. Bar plots show relative proportion of (tumor) cells. Data represent two independent tumors per genotype. **H.** Boxplots displaying the distribution of Hallmark MYC_TARGETS_V1 scores for snRNA-seq data from RPM and RPMM tumors. Statistical significance was assessed using a Wilcoxon rank-sum test. **I.** Spatial transcriptomic analysis of RPM and RPMM tumors using 10X Visium HD. Top panels show H&E-stained tumor sections used to define punch regions. Corresponding H&E images of the punched tumor areas are shown below. Spatial spots (dots), each representing a single cell, are colored by Hallmark MYC_TARGETS_V1 module score. **J.** Tumor cell vs. non-tumor cell composition of spatial transcriptomic profiles from RPM and RPMM tumors. Data represent three independent tumor punches per genotype. **K.** Relative cell cycle phase composition of spatial transcriptomic profiles from RPM and RPMM tumors. Data represent three independent tumor punches per genotype. **L.-M.** Trajectory inference was performed on scRNA-seq tumor cells using Monocle3. From the tumor cell compartment, RPM and RPMM tumor cells were randomly downsampled to equal cell numbers to account for unequal cell numbers between genotypes. Trajectories were learned with use_partition=FALSE to force construction of a single trajectory graph across all cells. Pseudotime was ordered using as root the cells with the top 5% highest CytoTRACE score. Separate trajectory analyses were then performed for RPM (L.) and RPMM (M.) separately.

Transcriptomic profiling (3’UTR sequencing of tumors, snRNA-seq of tumors and rRNA depleted RNA-seq of cell lines) and subsequent gene set enrichment analysis (GSEA) uncovered pronounced transcriptional differences between RPM and RPMM models (Figure 1D, Supplementary Figure 1E, F, Supplementary Table 1). In particular, both Myc Hallmarks (MYC_TARGETS_V1 and MYC_TARGETS_V2) showed enrichment in RPMM tumors compared to RPM tumors. Consistent with enhanced Myc pathway activity, RPMM tumors also exhibited significant upregulation of cellular stress-related pathways, such as the Hallmarks UNFOLDED_PROTEIN_RESPONSE and HYPOXIA, compatible with an increase in cellular stress (Figure 1D).

Furthermore, we found a significant downregulation of the Hallmark PANCREAS_BETA_CELLS gene set, which encompasses key neuroendocrine lineage markers such as Chga, Insm1, NeuroD1, and Sst (Figure 1D, Supplementary Figure 1E). In line with this observation, analyses of the top 30 downregulated genes in RPMM tumors revealed a marked enrichment of NE-associated genes, including Nrxn1, Nrxn3, Snap25, and Homer2 (Supplementary Figure 1G).

To further dissect tumor heterogeneity, cellular states, and differentiation in situ, we performed snRNA-seq (Figure 1E-H, L and M) of RPM and RPMM tumors. The majority of cells were annotated as epithelial or mesenchymal cells (Figure 1E). More specifically, PNECs and pulmonary mesenchymal progenitor–like (PMP) cells, with a smaller subset annotated as secondary crest myofibroblasts (SCMFs) were distinguishable based on distinct transcriptional profiles consistent with their respective lineage identities (Figure 1F). PNECs and PMPs clustered together in the snRNA-seq UMAP, whereas SCMFs formed clusters with other mesenchymal cell populations, suggesting that SCMFs likely represent stromal rather than tumor cells (Figure 1F). Given the cell-of-origin of the model, we consider PNECs as bona fide tumor cells. To determine whether PMP cells likewise represent part of the malignant tumor compartment, we systematically compared PMP and PNEC populations across multiple orthogonal features. First, PNECs showed very high cell-type annotation confidence scores, whereas PMP cells displayed clearly lower annotation confidence (Supplementary Figure 1H). Both PMP and PNEC populations further exhibited elevated MYC target gene signature scores compared with healthy cells, including endothelial and immune populations (Supplementary Figure 1I). In addition, PNEC and PMP cells exhibited markedly increased proliferative activity compared to the healthy cell populations (Supplementary Figure 1J). We next assessed transcriptional differences between PNEC and PMP populations and found that the major distinguishing features were neuroendocrine lineage-associated gene programs (Supplementary Figure 1K, Supplementary Table 2). This observation is consistent with the enrichment of PMP cells in RPMM tumors, where neuroendocrine gene expression is globally reduced, indicating that the PNEC-PMP separation largely reflects a shift in neuroendocrine differentiation state rather than distinct tumor-extrinsic lineage identity. Accordingly, we defined both PMP and PNEC populations as tumor cells in subsequent analyses.

To further investigate the effects of Miz1 loss while preserving tissue architecture and spatial organization, we complemented the snRNA-seq analysis with spatial transcriptomics (Figure 1I). Using the same annotation strategy as applied to the snRNA-seq dataset, spatial transcriptomic spots were predominantly assigned to tumor cell identities in both RPM and RPMM tumors, consistent with the cellular composition identified by snRNA-seq (Figure 1J). Despite minimal differences in cellular annotation resolution between the spatial transcriptomics and snRNA-seq, both approaches revealed highly concordant biological alterations associated with loss of Miz1 DNA-binding function. Across spatial transcriptomics and snRNA-seq data, tumor cells in RPMM samples displayed a pronounced shift in cell cycle distribution, with a reduced fraction of G1-phase cells and a concomitant significant increase in cells in the G2/M phase, indicative of enhanced proliferative activity (Figure 1 G, K). Consistent with this, RPMM tumor cells exhibited significantly elevated Myc target gene expression scores in both datasets (Figure 1H, I, Supplementary Figure 1L), corroborating the strong Myc pathway activation observed in bulk RNA-seq.

To further assess intratumoral heterogeneity and differentiation dynamics, we performed pseudotime trajectory analysis on snRNA-seq data (Figure 1L, M). In both RPM (Figure 1L) and RPMM tumors (Figure 1M), cells were organized along continuous trajectories, reflecting gradual transcriptional transitions across tumor cell states. Notably, RPM tumors exhibited a more branched trajectory structure, suggesting greater diversification of cellular states, whereas RPMM tumors followed a more linear and constrained trajectory. Quantification of pseudotime state distribution using Pielou’s evenness revealed a significant reduction in RPMM tumors compared to RPM (Supplementary Figure 1M), indicating a less even occupancy of the inferred trajectory space. Combined with the reduced branching structure, these findings indicate decreased cellular state diversity in RPMM tumors.

Collectively, these findings indicate that loss of Miz1 DNA-binding function in RPM tumors promotes increased Myc signaling while suppressing neuroendocrine differentiation programs, resulting in reduced cellular heterogeneity, a more uniform and less differentiated tumor cell state, and slower tumor progression.

### Cooperative Loss of Miz1 and Myc on Chromatin Defines a Distinct Transcriptional Subprogram

To better understand the consequences of Miz1’s transcriptional function in RPMM vs RPM tumors, we performed a genome-wide comparison of DNA-binding occupancy for Miz1 and Myc between RPM and RPMM samples using the DynaTag method ^14^. Profiling of Miz1 revealed a widespread loss of Miz1 chromatin binding in RPMM tumors, with 26,647 genomic sites showing reduced Miz1 occupancy compared to RPM tumors, validating the novel RPMM model and confirming that deletion of the BTB/POZ domain effectively abrogates Miz1 DNA binding (Figure 2A). Strikingly, the Miz1^ΔPOZ^ mutation induced extensive remodeling of Myc chromatin occupancy in RPMM tumors, with 6,796 regions showing increased Myc binding and 6,450 regions showing reduced binding relative to RPM tumors (Figure 2A).

**Figure 2:**
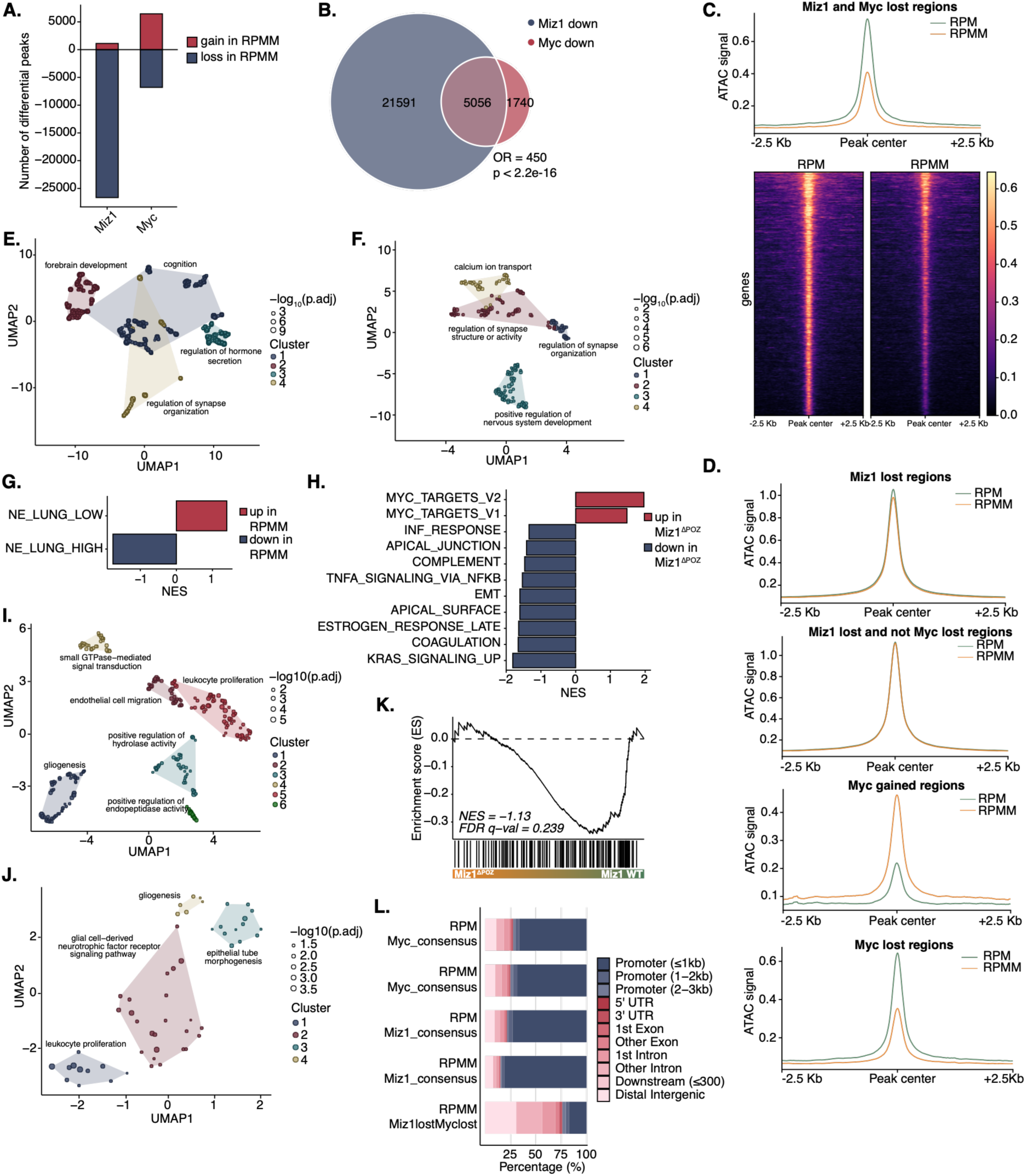
**A.** Differential DNA occupancy of Miz1 (left bar) and Myc (right bar) between RPM and RPMM across the genome. Positive bars (red) represent the number of peaks with increased occupancy in RPMM relative to RPM, whereas negative bars (blue) indicate the number of peaks with decreased occupancy in RPMM. **B.** Venn diagrams illustrating the overlap between genomic regions affected by Miz1 loss and Myc loss in RPMM samples. The overlap of peaks (400 bp width) with reduced Miz1 and Myc binding in RPMM is shown. Odds ratio (OR) and significance of overlap determined by Fisher’s exact test are indicated. **C.-D.** ATAC-seq signal intensity in RPM and RPMM tumors at Myc- and Miz1-associated genomic regions. Average profiles (C.-D.) and heatmaps (C.) show ATAC-seq signal centered on peak regions (±2.5 kb) for the indicated peak sets, comparing RPM and RPMM tumors. **E.** GO term enrichment of the overlapping genes that are regulated by loss of Miz1 and Myc in RPMM tumors. Significantly enriched GO terms were clustered by semantic similarity and shown in a UMAP plot. **F.** GO term enrichment of all significantly downregulated genes (log2FC<-1, FDR<0.05) in the RPMM model. 3’UTR bulk RNA-seq data of FFPE tumor tissue was used. Significantly enriched GO terms were clustered by semantic similarity and shown in a UMAP plot. **G.** GSEA of NE_LUNG_LOW and NE_LUNG HIGH gene sets from 3’UTR bulk RNA-seq of FFPE tissue isolated from RPMM and RPM tumors. **H.** GSEA of Hallmark pathways of bulk RNA-seq of DLBCL mouse tumors with Miz1^ΔPOZ^ and Miz1^WT^. Pathways with p-value < 0.05 were included. **I.** GO term enrichment of all significantly downregulated genes (log2FC<-1, FDR<0.05) in the Miz1ΔPOZ DLBCL model. Significantly enriched GO terms were clustered by semantic similarity and shown in a UMAP plot. **J.** GO term enrichment of 167 DLBCL-associated genes that are downregulated in Miz1^ΔPOZ^ B-cell lymphoma tumors. These genes were defined by intersecting genes preferentially expressed in B-cell versus T-cell lymphoma with genes downregulated in Miz1^ΔPOZ^ versus Miz1^WT^ B-cell lymphoma. Significantly enriched GO terms were clustered by semantic similarity and shown in a UMAP plot. **K.** GSEA of the 150 DLBCL driver genes using bulk RNA-seq of B-ALL/lymphoma tumors with Miz1^ΔPOZ^ vs Miz1^WT^ expression. **L.** Genomic distribution of Myc and Miz1 consensus peaks in RPM and RPMM tumors, as well as peaks with cooperative loss of Myc and Miz1 in RPMM tumors, categorized by promoter and distal regions.

To assess whether loss of Miz1 and Myc binding occurs at shared genomic regions in RPMM tumors, we intersected Miz1- and Myc-depleted peaks. This analysis revealed 5,056 overlapping regions, representing ∼75% of all sites with reduced Myc binding in RPMM tumors, indicating a frequent cooperative loss of Miz1 and Myc occupancy at the same genomic loci (Figure 2B). To determine whether the concurrent loss of Miz1 and Myc binding was associated with changes in chromatin accessibility, we integrated Assay for Transposase-Accessible Chromatin (ATAC) sequencing (ATAC-seq) data from RPM and RPMM tumors. At genomic regions with cooperative loss of Miz1 and Myc binding, ATAC-seq profiles revealed a pronounced reduction of chromatin accessibility in RPMM compared to RPM tumors, indicating chromatin closure at these sites (Figure 2C). Loss of Miz1 binding alone did not measurably affect chromatin accessibility, whereas changes in Myc occupancy were accompanied by corresponding changes in ATAC signal (Figure 2D, Supplementary Figure 2A–D). Importantly, genome-wide ATAC-seq profiles were highly similar between RPM and RPMM tumors, with near-identical overall chromatin accessibility and strong correlation of ATAC signal across samples (Supplementary Figure 2E), indicating that these accessibility changes are locus-specific rather than global, and specifically associated with alterations in Myc occupancy (Supplementary Figure 2C-D).

Gene ontology (GO) term enrichment analysis and subsequent semantic similarity clustering of the genes regulated by the cooperatively Miz1- and Myc-lost regions revealed a strong link to neural and hormonal processes, including regulation of neurogenesis, and hormone transport and secretion (Figure 2E, Supplementary Table 2). Consistent with this, GO term analyses revealed that genes significantly downregulated in RPMM tumors were similarly enriched for neuroendocrine-related gene sets (Figure 2F, Supplementary Figure 2F, G, Supplementary Table 2). To directly assess the impact on neuroendocrine identity, we performed GSEA using two predefined 25-gene SCLC signatures from Zhang et. al ^32^. This analysis demonstrated a significant decrease in the NE_LUNG_HIGH signature - genes associated with high neuroendocrine differentiation in SCLC - and a concomitant increase in the NE_LUNG_LOW signature - genes associated with low neuroendocrine differentiation in SCLC - in RPMM tumors (Figure 2G, Supplementary Table 1). In summary, these analyses indicate a cooperatively Miz1- and Myc-regulated program in SCLC that is strongly lineage-specific, converging on neuroendocrine differentiation characteristics, a hallmark of SCLC.

To further specify the effects of MIZ1 in MYC-driven cancer, we analyzed an independent dataset from a Myc-driven diffuse large B-cell lymphoma (DLBCL) tumor model comparing Miz1^WT^ and Miz1^ΔPOZ20.^ Also in this setting, Miz1^ΔPOZ^ tumors showed increased expression of Myc target gene sets, consistent with conserved Myc pathway activation upon loss of Miz1 DNA-binding function (Figure 2 H, Supplementary Table 1). Strikingly, GSEA showed a significant decrease in the Hallmark gene set COMPLEMENT. This gene set is enriched for complement components and regulators that are highly expressed in germinal center and activated B cells, where complement receptor signaling supports B-cell survival and antigen-driven responses ^33^. In line with this, GO term enrichment of genes downregulated in DLBCL Miz1^ΔPOZ^ tumors highlighted B cell lineage-associated processes, including leukocyte proliferation (Figure 2I, Supplementary Table 2). To assess whether the genes downregulated upon Miz1^ΔPOZ^ mutation reflected loss of B-cell lineage-associated programs, we intersected genes preferentially expressed in B-cell compared with T-cell lymphoma with genes downregulated in Miz1^ΔPOZ^ versus Miz1^WT^ B-cell lymphoma tumors. This identified 167 B-cell lineage-associated genes lost upon Miz1^ΔPOZ^ mutation, indicating that a large fraction of downregulated genes is specific to a B cell lymphoma context rather than broadly shared across lymphoid malignancies. GO term enrichment analysis of these 167 genes identified leukocyte proliferation as a dominant functional cluster, along with glial cell-derived neurotrophic factor receptor signaling - both pathways with established relevance in B-cell malignancies (Figure 2J, Supplementary Table 2)^34^. Consistent with a lineage-specific transcriptional impact, GSEA using a curated set of 150 DLBCL driver genes ^35^ showed a trend toward reduced expression of these genes in Miz1^ΔPOZ^ tumors, although this did not reach statistical significance (Figure 2K). Together, these findings indicate that loss of chromosomal binding of Miz1 may universally activate Myc target programs, while its impact on lineage-specific transcriptional programs may vary between entities.

Finally, we examined the genomic distribution of Miz1 and Myc binding sites. The vast majority of Miz1 and Myc peaks in RPM and RPMM tumors (∼75-85%) were localized to promoter regions, with minimal differences between the genotypes. However, in stark contrast, the genomic regions that exhibited concurrent loss of Miz1 and Myc binding in RPMM tumors showed a striking redistribution away from promoters to distal regulatory elements. Only ∼25% of these shared Miz1/Myc binding depleted sites were found at promoters, while the remaining ∼75% were located at distal, putative enhancer elements (Figure 2L).

Thus, loss of Miz1 and Myc occupancy in RPMM relative to RPM tumors preferentially co-occurs at distal regulatory elements rather than promoters and is associated with reduced expression of neuroendocrine gene programs. These findings point to a potentially enhancer related relationship between Miz1 and Myc in RPM tumors, distinct from their previously described antagonistic promoter-based interactions ^16^.

### Cooperative Binding of Miz1 and Myc at Enhancers Maintains Neuroendocrine Differentiation

Next, we sought to characterize the transcriptional impact of regions showing shared loss of Miz1 and Myc binding in RPMM tumors. Because the majority of these regions fall outside of promoter zones, we employed the Activity-By-Contact (ABC) model ^27^ to predict their putative target genes based on enhancer-promoter interactions. From a total of 5,056 Miz1/Myc co-bound regions lost in RPMM, the ABC model predicted 9,729 enhancer-promoter loops, corresponding to 6,790 unique target genes (Supplementary Table 3). GSEA using the enhancer target gene set showed significant downregulation in the RPMM tumors (Figure 3A). Furthermore, GO term enrichment analysis of this full enhancer target set revealed strong associations with neuroendocrine signaling pathways (Supplementary Figure 3A, Supplementary Table 2). Similar enrichment patterns were observed across additional datasets, including RPMM cell lines analyzed by 3ʹUTR bulk RNA-seq (Supplementary Figure 3B), spatial transcriptomics of RPMM tumors (Supplementary Figure 3C), and snRNA-seq data (Supplementary Figure 3D).

**Figure 3:**
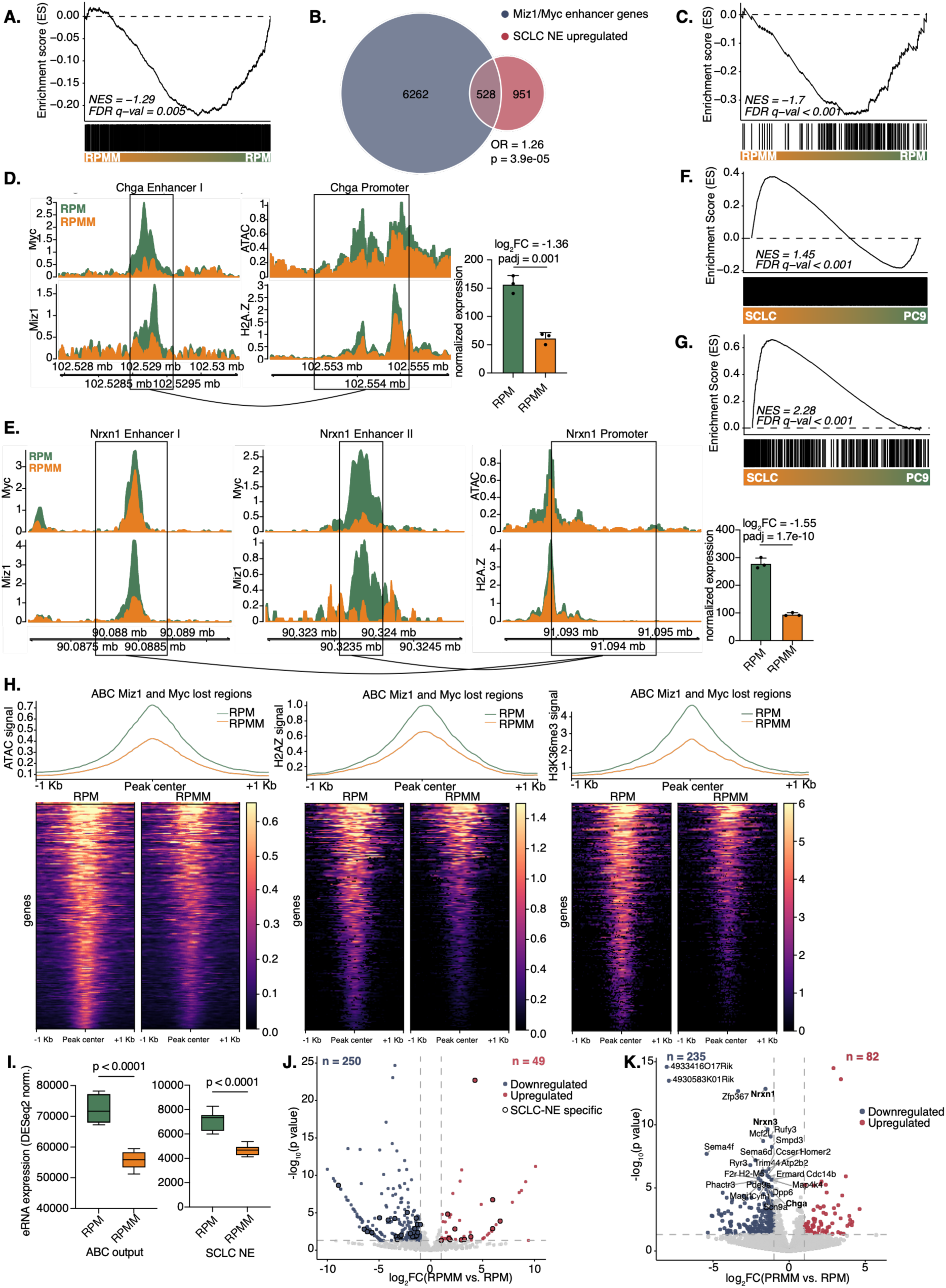
**A.** GSEA of the 6,790 unique target genes of the ABC output using 3’UTR bulk RNA-seq of FFPE tissue isolated from RPMM and RPM tumors. **B.** Venn diagram showing the overlap of genes regulated by cooperative Miz1 and Myc binding loss in RPMM and neuroendocrine genes being upregulated in SCLC versus other cancer entities. Odds ratio (OR) and significance of overlap determined by Fisher’s exact test are indicated. **C.** GSEA of the 528 unique target genes of the ABC output overlapping the SCLC specific neuroendocrine genes using 3’UTR bulk RNA-seq of FFPE tissue isolated from RPMM and RPM tumors. **D.-E.** Signal intensity of Miz1 and Myc (left) as well as ATAC and H2A.Z (right) in RPM (green) and RPMM (orange) tumors. The ABC-predicted enhancer and promoter region of Chga (D.) and Nrxn1 (E.) are shown. The bar plot represents the normalized RNA expression of the respective gene using 3’UTR bulk RNA-seq of FFPE tissue isolated from RPMM and RPM tumors. Log2FC and padj were obtained from the DESeq2 differential expression analysis. **F.-G.** GSEA of the 6,790 unique ABC model target genes (J.) or the 528 unique target genes of the ABC output overlapping with the SCLC specific neuroendocrine genes (K.) from the murine model, mapped to human orthologs, using 3’UTR bulk RNA-seq from human SCLC cell lines (SCLC21H and DMS273) compared to the human NSCLC cell line PC9. **H.** Signal intensity of ATAC-seq as well as H3K36me3 and H2AZ DynaTag of RPM and RPMM tumors at ABC enhancer regions. Self-promoter regions were excluded. **I.** eRNA expression at Miz1- and Myc-bound enhancer regions (ABC_output) and the further filtered ABC_output regions overlapping with SCLC specific neuroendocrine enhancer regions (SCLC NE, right) of 3’UTR bulk RNA-seq of FFPE tissue isolated from RPMM and RPM tumors is shown. Self-promoter regions were excluded. T-test was used to compare RPM with RPMM tumors. **J.** Volcano plot of eRNAs at Miz1 and Myc co-bound enhancer regions based on 4sU RNA-seq data comparing RPMM575 vs RPM426. Self-promoter regions were excluded. Significantly downregulated enhancer RNAs (log₂FC < –1, p < 0.05) are shown in blue, significantly upregulated ones (log₂FC > 1, p < 0.05) in red. SCLC NE-specific enhancer regions are highlighted with a black outline. **K.** Volcano plot of genes regulated by cooperative Miz1 and Myc binding. Gene expression of 3’UTR RNA-seq of FFPE tissue isolated from RPMM and RPM tumors is shown. Significantly downregulated (log2FC<-1, p<0.05) gene expression of Miz1 and Myc enhancer regulated genes are depicted in blue and significantly upregulated (log2FC>1, p<0.05) gene expression are depicted in red.

To investigate whether Miz1/Myc-bound enhancers in RPM tumors are linked to SCLC associated neuroendocrine gene regulatory programs, we performed a multi-step integrative analysis. For this, we used the 6,790 unique genes predicted by the ABC model to be targets of enhancers co-bound by Miz1 and Myc in RPM tumors. To identify biologically enriched categories within this gene set, we compared it to genes associated with hallmark features of SCLC. In detail, we used a dataset comparing gene expression across SCLC versus 33 distinct cancer entities ^36^. From this dataset, we selected all genes significantly upregulated in SCLC (log2FC > 1) compared to non-SCLC tumors. We then performed GO term enrichment on these upregulated genes and selected the top 50 enriched biological processes associated with neuroendocrine features. From these pathways, we extracted the genes contributing to the enrichment in SCLC, resulting in a curated list of 1,479 genes highly expressed in SCLC and linked to functionally enriched neuroendocrine biological processes. Intersecting this SCLC-specific, NE-associated gene set with the Miz1/Myc co-regulated enhancer targets yielded 528 overlapping genes, representing a substantial convergence (∼36% of all SCLC-specific neuroendocrine associated genes) and suggesting that enhancer co-binding by Miz1 and Myc may be involved in activating a transcriptional program highly characteristic of SCLC (Figure 3B). To determine whether this enhancer-driven transcriptional program actually altered gene expression in the RPMM tumors, we performed GSEA on RNA-seq data from RPMM and RPM tumors using the 528 gene set. This revealed a strong and statistically significant downregulation of Miz/Myc enhancer regulated neuroendocrine genes in RPMM (Figure 3C). In contrast, a similar GSEA using the full set of 1,479 SCLC-upregulated neuroendocrine genes showed only a non-significant trend toward downregulation in RPMM tumors (Supplementary Figure 3E).

To assess the functional consequences of enhancer-associated Miz1/Myc binding at key neuroendocrine loci, we examined 4 representative canonical neuroendocrine genes - Chga (Figure 3D), Nrxn1 (Figure 3E), Nrxn3, and Grin3a (Supplementary Figure 3F) - that were predicted by the ABC model to be regulated by Miz1/Myc-bound enhancers. At predicted enhancer regions, Miz1 and Myc occupancy was strongly reduced in RPMM compared to RPM tumors. This loss of enhancer binding was accompanied by decreased chromatin accessibility at the corresponding promoters, as reflected by reduced ATAC-seq signal and diminished H2A.Z deposition, indicative of reduced promoter activity. Consistent with these chromatin changes, RNA expression of the respective genes was significantly reduced in RPMM relative to RPM tumors, directly linking loss of Miz1/Myc enhancer binding to promoter chromatin closure and transcriptional downregulation. To assess whether Miz1/Myc co-regulated enhancer regions form higher-order regulatory units, we performed an analysis for super-enhancers on ABC-defined enhancer regions in RPM tumors using the ROSE algorithm ^37,38^. Enhancers were stitched (within 12.5 kb, excluding ±2 kb around transcription start sites) and ranked by ATAC-seq signal. While a subset of highly accessible enhancer clusters met criteria for super-enhancers, these regions did not regulate a distinct subset of target genes, suggesting that Miz1/Myc-dependent transcription is driven by distributed enhancer activity rather than discrete super-enhancer loci (Supplementary Figure 3G, Supplementary Table 4).

To assess the relevance of the MIZ1/MYC enhancer program in human SCLC, we performed a cell line-specific integrative analysis using ATAC-seq, MYC ChIP-seq, and RNA-seq data from MYC-amplified human SCLC cell lines ^39^. For each cell line, a separate ABC model (Supplementary Table 3) was generated using the corresponding ATAC-seq data to predict enhancer-promoter interactions in a human, cell line-specific context. We then restricted the analysis to the same set of 6,790 unique target genes that were predicted by the murine ABC model to be regulated by Miz1/Myc co-bound enhancers lost in RPMM tumors. For these genes, we identified the corresponding enhancer regions in the human ABC model and stratified them based on MYC binding using MYC ChIP-seq data. Gene expression levels were subsequently compared between genes linked to MYC-bound enhancers and those linked to enhancers lacking MYC binding. Across three MYC-amplified human SCLC cell lines, genes associated with MYC-bound enhancers exhibited significantly higher expression than non-MYC-bound counterparts (Supplementary Figure 3H), supporting a conserved role of enhancer-associated MYC binding in promoting transcriptional output at MIZ1/MYC-regulated loci across murine and human SCLC models.

To further extend our analyses to human patient-derived models, we next compared gene expression between human SCLC and NSCLC cell lines using 3’UTR RNA-seq. As expected, genes associated with SCLC-specific neuroendocrine signaling showed higher expression in SCLC compared to NSCLC (Supplementary Figure 3I), consistent with the known lineage-specific transcriptional program. We then asked whether genes linked to MIZ1/MYC co-bound enhancers in the murine model exhibit a similar pattern in human cells. Notably, the full set of ABC model target genes was significantly enriched in SCLC relative to NSCLC (Figure 3F). This difference became even more pronounced when focusing on the subset of ABC target genes overlapping SCLC-specific neuroendocrine genes (Figure 3G), indicating a strong convergence between enhancer-associated MIZ1/MYC regulation and lineage-defining transcriptional programs.

We next questioned whether Miz1 and Myc co-binding is associated with enhancer activation. To this end, we focused on enhancer regions identified by the ABC model that show cooperative binding of Miz1 and Myc in RPM tumors, but loss of both transcription factors in RPMM. To ensure we were evaluating bona fide distal regulatory elements, we excluded all ABC enhancers that overlapped annotated promoter regions, regardless of the target gene. We examined chromatin accessibility at these enhancer sites using ATAC-seq. In RPM tumors, these regions displayed high accessibility, consistent with an active enhancer state. In contrast, accessibility was markedly reduced in RPMM tumors, suggesting that Miz1 and Myc binding in the RPM model is associated with an open, active chromatin state, which becomes impaired upon their loss in the RPMM model (Figure 3H, left graphic). Characterizing the chromatin environment at these sites, we analyzed the histone variant H2A.Z, a marker of nucleosome remodeling at regulatory elements, and H3K36me3, which is associated with transcriptional elongation^40,41^. Both marks were enriched at Miz1/Myc-bound enhancers in RPM but substantially diminished in RPMM (Figure 3H, middle and right graphic). This coordinated loss of accessibility, H2A.Z deposition, and elongation-associated histone marking suggests that Miz1 and Myc co-binding promotes a transcriptionally active enhancer configuration in RPM that is epigenetically silenced following their loss in RPMM.

To further assess whether cooperatively Miz1- and Myc-bound enhancers in RPM tumors are transcriptionally active, we examined enhancer RNA (eRNA) expression at these regions. Using 3ʹUTR bulk RNA-seq data from RPM and RPMM tumors, we quantified eRNA levels at both the complete set of ABC-predicted Miz1/Myc-bound enhancers (ABC output; Figure 3I, left) and the more focused 528 genes regulated by Miz1 and Myc with SCLC specific neuroendocrine features (SCLC NE; Figure 3I, right). This analysis revealed a significant reduction in eRNA expression at the predefined regions in RPMM tumors compared to RPM. To more efficiently capture nascent and unstable eRNAs, we performed 4sU-sequencing, which selectively enriches for newly transcribed RNAs ^42^. Of the predicted ABC target genes, 250 eRNAs were significantly lower expressed in RPMM tumors, whereas only 49 showed increased expression (Figure 3J). This marked imbalance further supports the idea that co-binding of Miz1 and Myc in RPM tumors is essential for maintaining active enhancer transcription, particularly at NE-specific regulatory regions, a function that is broadly impaired upon their loss in RPMM tumors.

To further validate these findings, we examined individual gene expression changes in our 3’UTR bulk RNA-seq dataset of all enhancer-regulated genes identified by the ABC model. A differential expression analysis revealed a clear bias toward downregulation in RPMM tumors, with 235 genes significantly downregulated and only 82 significantly upregulated (Figure 3K). This imbalance further supports the notion that Miz1 and Myc co-binding at enhancer regions predominantly drives transcriptional activation. Several core neuroendocrine genes were significantly downregulated, including Chga, Nrxn1, and Nrxn3, highlighting the functional impact of enhancer-mediated regulation. These findings underscore the relevance of the cooperative Myc/Miz1 enhancer binding to directly modify lineage-defining genes in murine SCLC models and human SCLC cells.

### Miz1 facilitates Myc binding to low-affinity E-boxes at distal regulatory elements

To further define the regulatory potential of the genomic regions that showed a cooperative loss of Miz1 and Myc binding in RPMM tumors, we performed motif enrichment analysis using HOMER. This analysis revealed a striking overrepresentation of DNA motifs recognized by transcription factors involved in neuroendocrine differentiation, including Ascl1 (Figure 4A), Atoh1, NeuroG1, and NeuroD1 (Supplementary Figure 4A). Analysis of the enriched motifs revealed that the vast majority of significantly overrepresented sequences contained the core CAGCTG motif, an E-box with a lower MYC affinity compared to the high affinity E-box CACGTG, and thus typically requiring cooperative interactions for stable MYC binding ^43^ (Figure 4A, Supplementary Figure 4A). Because Myc binding at these regions is selectively lost in the absence of DNA-bound Miz1, this finding led us to hypothesize that Miz1 facilitates Myc recruitment to low-affinity E-boxes, rather than acting through independent or redundant binding mechanisms.

**Figure 4:**
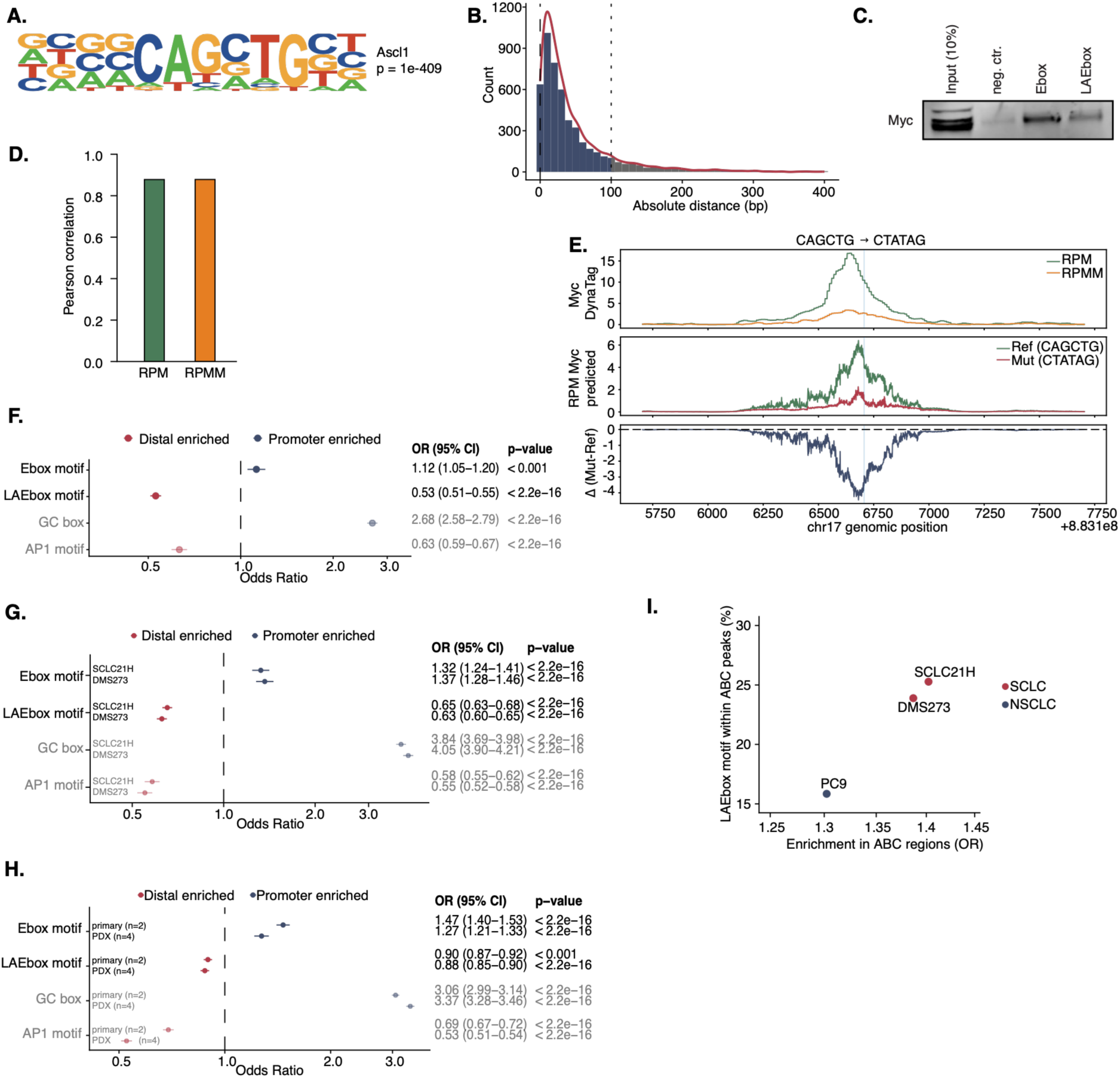
**A.** HOMER motif enrichment analysis of Miz1 and Myc lost regions in RPMM tumors, showing the second top enriched motif. **B.** Distribution of absolute distances between Miz1 and Myc peak center within Miz1 and Myc lost regions in RPMM tumors. **C.** Western blot analysis of DNA pulldown assays. Myc binding to the canonical E-box (CACGTG) and the low-affinity E-box (CAGCTG) using RPM150 cells. **D.** Performance of the fine-tuned AlphaGenome model on independent test regions. Pearson correlations between observed and predicted Myc DynaTag signals are shown for RPM and RPMM samples. **E.** In silico mutagenesis using the fine-tuned AlphaGenome model. Mutation of the candidate Myc motif (CAGCTG) to a scrambled sequence (CTATAG) is shown. Top, observed DynaTag signal; middle, predicted signal for reference and mutant sequences; bottom, change in predicted signal (Δ mutant − reference). **F.-H.** ATAC-seq peaks in RPM tumors (F.), hSCLC and NSCLC cell lines (SCLC21H, DMS273, PC9; G.), or FFPE SCLC patient biopsy (primary) or an SCLC PDX sample (PDX; H.) containing the E-box motif (CACGTG), the low-affinity E-box motif (LAEbox, CAGCTG), the GC box (positive control for promoter enriched motif, RSGVGS) or the AP1 motif (positive control for distal enriched motif, TGABYCA). Peaks were stratified by distal (all non-promoter regions) and promoter regions (TSS ± 3kb). Forest plot showing odds ratios for motif enrichment in distal versus promoter peaks. Error bars indicate 95% confidence intervals. Statistical significance was assessed using Fisher’s exact test. **I.** Enrichment of the low-affinity E-box (LAEbox) motif within ABC-defined enhancer regions in human SCLC and NSCLC cell lines. Scatter plot showing the odds ratio for enrichment of LAEbox motifs in ABC regions compared to ATAC-seq peaks outside of ABC-defined interactions (x-axis) and the fraction of ABC regions containing the LAEbox motif (y-axis).

Because the overlapping Miz1–Myc lost regions were defined using fixed 400 bp peak widths—where overlap can occur even if only a single base pair is shared—we next quantified the absolute distance between the centers of Miz1 and Myc peaks within these regions. Strikingly, we found that ∼88% of Miz1 and Myc peaks were located within 100 bp of each other (Figure 4B). This close spatial proximity provides further support for a cooperative binding model, rather than independent recruitment events.

Based on this observation, we hypothesized that Miz1 binds a nearby DNA motif and thereby facilitates recruitment of Myc to the low-affinity E-box.

Guided by these in silico findings, we performed biochemical DNA pulldown assays to experimentally validate the binding specificities of Myc. Consistent with previous reports, Myc bound robustly and specifically to the canonical E-box (CACGTG) and, to a lesser extent, to the low-affinity E-box (CAGCTG) (Figure 4C, Supplementary Figure 4B).

To independently evaluate the functional relevance of the identified motif in their native genomic context, we trained AlphaGenome ^28^ on our DynaTag datasets for Myc binding in RPM and RPMM tumors. The models accurately predicted Myc occupancy in held-out test regions, achieving high Pearson correlations for RPM_Myc (0.88) and RPMM_Myc (0.88) signal tracks (Figure 4D, Supplementary Table 5).

We next performed *in silico* mutagenesis of the low-affinity E-box (CAGCTG) within the Miz1- and Myc-lost regions. Mutation of the motif to a scrambled sequence (CTATAG) resulted in a pronounced decrease in predicted Myc binding (Figure 4E, Supplementary Table 5). Notably, 96% of mutated sites showed reduced Myc signal in the RPM model, demonstrating a highly consistent dependence of Myc occupancy on the low-affinity E-box motif at these regions.

We next asked why this cooperative binding configuration preferentially occurs at enhancer regions. To address this, we examined the genomic distribution of different motifs within all accessible chromatin regions in RPM defined by ATAC-seq, including the canonical E-box (Ebox motif) and low-affinity E-box (LAEbox motif) (Figure 4F). As expected, the canonical E-box motif was enriched at promoter regions. In contrast, the low-affinity E-box showed a clear enrichment at distal regulatory elements. In line with established motif distributions, the GC-box, a promoter-associated element, was strongly enriched at promoters, whereas the AP-1 motif, a canonical enhancer-associated motif, was enriched at distal regions (Figure 4F). This enrichment pattern provides a mechanistic explanation for the preferential loss of Miz1 and Myc binding observed at enhancers in RPMM tumors.

Importantly, this distribution was not restricted to the murine RPM model but was conserved across human systems. Performing the same analysis in human SCLC cell lines (Figure 4G) as well as in patient-derived material, including primary biopsies as FFPE samples and PDX models (Figure 4H), revealed a highly consistent pattern: canonical E-box motifs remained preferentially enriched at promoters, whereas the low-affinity E-box was significantly enriched at distal regulatory elements. Together, these findings demonstrate that the motif architecture underlying cooperative binding of MIZ1 and MYC at enhancers is conserved between mouse models and human SCLC, reinforcing its biological relevance.

To further assess the relevance of this enhancer-associated motif architecture in a human, lineage-specific context, we performed an integrative analysis using human SCLC and NSCLC cell lines. Given that SCLC is characterized by a neuroendocrine transcriptional program (see Figure 3J,K), we reasoned that NSCLC would serve as an appropriate negative control for lineage-specific regulatory features. For each cell line, we generated cell line-specific ABC model based on ATAC-seq and H3K27ac data to predict enhancer-promoter interactions (Supplementary Table 3). Notably, ATAC-seq and H3K27ac profiles were largely comparable between SCLC and NSCLC at promoter-proximal regions, while distal regulatory elements displayed pronounced lineage-specific differences (Supplementary Figure 4C), suggesting that enhancer landscapes rather than promoter architecture distinguish the neuroendocrine SCLC state. We then focused on the set of genes identified by the murine ABC model as targets of MIZ1/MYC co-bound enhancers lost in RPMM tumors and assessed whether corresponding enhancer–promoter interactions could be recovered in the human ABC models. We examined the distribution of the low-affinity E-box motif (LAEbox), the preferential MYC binding motif in presence of MIZ1, within ABC-defined enhancer regions. Strikingly, the fraction of ABC regions containing low-affinity E-box motifs was markedly higher in MYC-amplified human SCLC cell lines (SCLC21H and DMS273) compared to the NSCLC cell line PC9 (Figure 4I), highlighting a lineage-specific enrichment of this motif in SCLC regulatory elements. Moreover, within SCLC models, the low-affinity E-box motifs were preferentially enriched in ABC regions compared to accessible chromatin regions outside of predicted enhancer-promoter interactions, indicating that these motifs are specifically associated with functional enhancers (Figure 4I). Notably, this pattern was recapitulated in patient-derived and xenograft-based models, where both the primary FFPE sample and the corresponding PDX model likewise showed a high fraction of low-affinity E-box motifs within ABC-defined enhancer regions (Supplementary Figure 4D).

These results support a model in which low-affinity E-box-driven regulatory elements contribute to a lineage-specific enhancer program in SCLC, as evidenced by their preferential enrichment within ABC-defined enhancer regions and their reduced presence in NSCLC. Collectively, our findings identify a composite MIZ1-MYC mechanism that enables cooperative binding at enhancer regions. This mechanism provides a molecular explanation for the enhancer-specific loss of MYC binding observed upon disruption of MIZ1 DNA engagement in RPMM tumors and establishes a conserved, lineage-restricted regulatory principle underlying MYC function in neuroendocrine SCLC. Importantly, the regulatory architecture is not restricted to the murine model but is consistently observed across human SCLC cell lines and patient-derived samples, underscoring its conserved biological relevance.

### Promoter-Localized Loss of Miz1 Relieves Repression of Myc-Driven Transcriptional Programs and Increases Chemotherapeutic Sensitivity

To better understand how the Miz1^ΔPOZ^ mutation impacts transcriptional regulation, we next extended our analysis to a genome-wide scale. DynaTag profiling revealed that the Miz1^ΔPOZ^ mutation not only results in regions of reduced Myc binding but also leads to the emergence of genomic regions with increased Myc occupancy (see Figure 2A). Importantly, the number of gained Myc peaks was comparable to the number of lost peaks, suggesting that Myc binding is not globally changed, but rather redistributed across the genome following loss of DNA-bound Miz1. In line with this, Myc RNA expression itself remained unchanged between RPM and RPMM tumors (log2FC = 0.56, padj = 0.59), indicating a model in which loss of promoter-bound Miz1, an established negative regulator of Myc-dependent transcription ^45^, together with increased Myc binding, shifts the regulatory balance toward transcriptional activation at Myc-bound promoters.

To determine whether regions of Miz1 loss and Myc gain converge on common regulatory targets, we intersected regions showing reduced Miz1 binding with those displaying increased Myc occupancy. Although the absolute overlap at the peak level was minimal (Figure 5A), gene-level mapping of these peaks to their nearest target genes revealed a substantial intersection: 2,229 genes were both associated with Miz1 loss and Myc gain in RPMM tumors, representing ∼65% of all genes linked to increased Myc binding in RPMM (Figure 5B). This spatial separation, with Miz1 loss predominantly at distal enhancers and Myc gain at promoter-proximal regions, is consistent with the broader genomic redistribution of Myc occupancy described above and suggests that the same gene loci are simultaneously subject to loss of enhancer-associated Miz1/Myc activity and relief of Miz1-mediated repression at their promoters.

**Figure 5:**
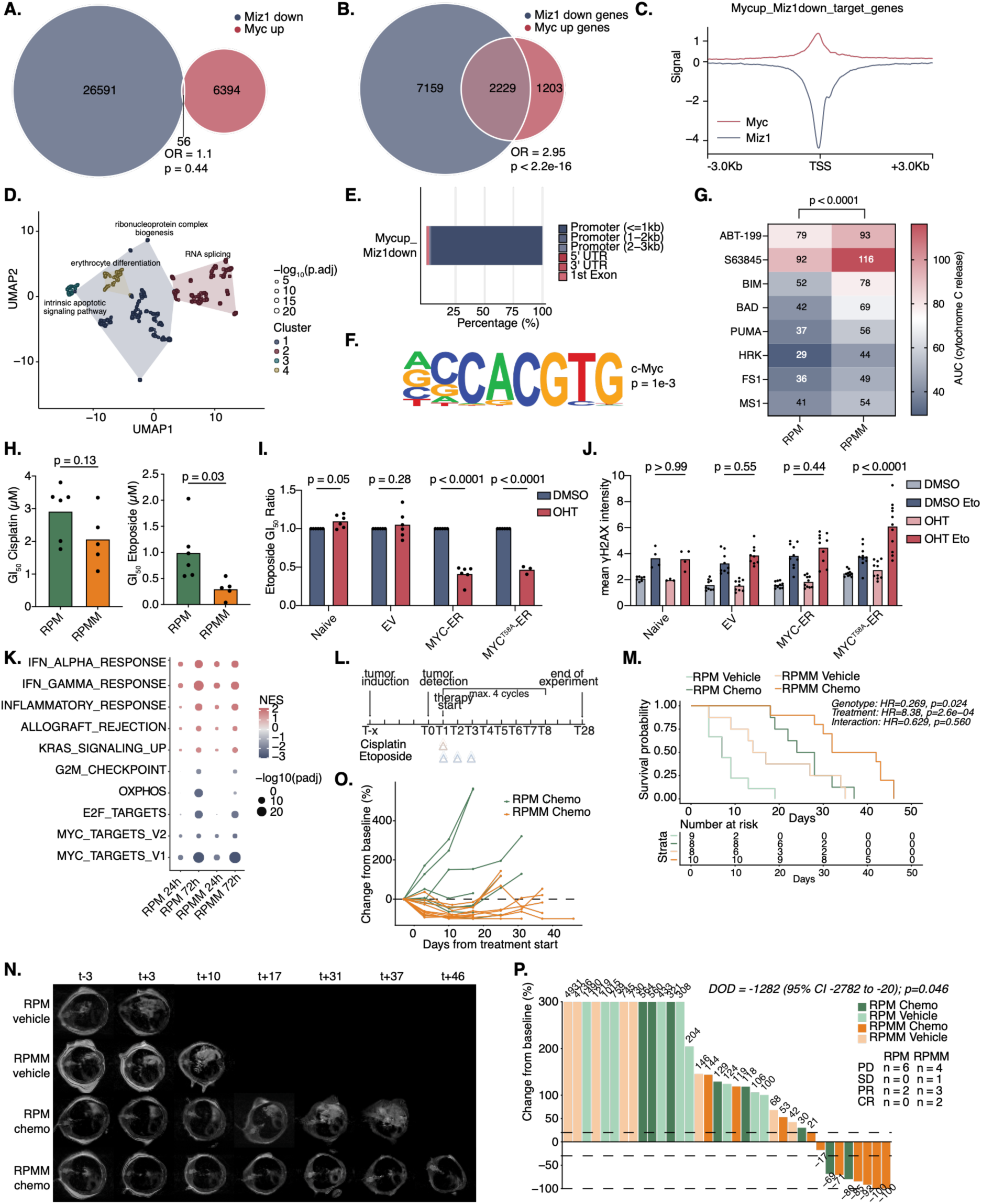
**A.-B.** Venn diagrams illustrating the overlap between regions and genes affected by Miz1 loss and Myc gain in RPMM samples. The overlap of peaks with reduced Miz1 binding and increased Myc binding in RPMM is shown (A., peak width=400bp). In the right diagram, each peak was assigned to its nearest gene, and the overlap of genes between reduced Miz1 binding and increased Myc binding is depicted (B.). Odds ratio (OR) and significance of overlap determined by Fisher’s exact test are indicated. **C.** Myc and Miz1 binding profile from DynaTag data on the transcription start site (TSS) of the 2229 genes associated with loss of Miz1 and gain of Myc. Differential binding values represent the comparison between RPMM and RPM occupancy, with negative values indicating reduced binding in RPMM relative to RPM, and positive values indicating enhanced binding in RPMM relative to RPM. **D.** GO term enrichment of the overlapping genes that are regulated by loss of Miz1 and gain of Myc in RPMM tumors. Significantly enriched GO terms were clustered by semantic similarity and shown in a UMAP plot. **E.** Genomic distribution of peaks with cooperative loss of Myc and gain Miz1 in RPMM tumors, categorized by promoter and distal regions. **F.** HOMER motif enrichment analysis of Miz1 loss and Myc gained regions in RPMM tumors, showing the canonical c-Myc binding motif. **G.** Heatmap of BH3 profiling showing sensitivity of RPM (n=6) and RPMM (n=5) cell lines against specific apoptosis-inducing peptides. Three biological replicates are included. Two-way ANOVA was used to compare the apoptotic priming sensitivity between RPM and RPMM. **H.** Cell viability assessed by CTG assay of RPM (n=6) and RPMM (n=5) cell lines after treatment with cisplatin (left) or etoposide (right). Three biological replicates are included. T-test was used to calculate the difference between RPM and RPMM cell lines. **I.-J.** Functional characterization of stably transduced RP1380 SCLC cells stably expressing empty vector (EV), MYC-ER, or MYCT58A-ER. Cell viability assessed by CTG assay, shown as the Etoposide GI50 Ratio (4-OHT treated compared to DMSO treated cells; I.). Mean γH2AX fluorescence intensity measured by immunofluorescence staining following the indicated treatments (J.). Two-way ANOVA was used to calculate significance. **K.** GSEA of etoposide-induced transcriptional changes in RPM and RPMM cell lines based on 3ʹUTR RNA-seq data. Normalized enrichment scores (NES) are shown for RPM and RPMM cells treated with etoposide for 24 h or 72 h relative to DMSO controls. Top 5 enriched and decreased hallmark gene sets are shown. Dot color indicates NES, dot size represents −log₁₀(padj), and a black outline denotes statistically significant enrichment (padj < 0.05). **L.** Schematic overview of treatment regimen. Mice inhaled CGRP-Cre virus. On the day after tumor detection, therapy with max. 4 cycles of cisplatin (5 mg/kg on day 1) and etoposide (10 mg/kg on day 1, 2, and 3) was started. Experiment was ended max. 28 days after therapy start. **M.** Kaplan-Meier Analysis of mice bearing RPM and RPMM tumors treated with either vehicle or chemotherapy (cisplatin + etoposide). Time after start of chemotherapy is shown. Statistical significance was assessed using a Cox proportional hazards model with an interaction term to test whether the survival benefit of chemotherapy differed between the genotypes (RPM and RPMM). **N.** Representative MR images of RPM and RPMM mice treated with either vehicle or chemotherapy. The representative images were acquired at the indicated time points (t, days after tumor induction). **O.** Spider plot of RPM and RPMM mice treated with chemotherapy showing the change of tumor size over time. **P.** Waterfall plot of RPM and RPMM tumor size at the endpoint of the experiment. The change from baseline (tumor size at treatment start) is shown. Statistical testing was performed using a bootstrap difference-in-differences approach (10,000 resamples) comparing the chemotherapy effect (Chemo–Vehicle) between RPM and RPMM. The reported DOD is the difference of treatment effects between genotypes, with 95% bootstrap confidence interval and two-sided corrected bootstrap p-value.

Aggregate profiling across transcription start sites of the overlapping 2229 genes co-regulated by Miz1 loss and Myc gain revealed a pronounced loss of Miz1 occupancy in RPMM tumors, accompanied by increased Myc binding at these loci (Figure 5C). GO term enrichment analysis revealed a pronounced enrichment in fundamental cellular processes, including ribosome biogenesis, RNA processing, and apoptosis (Figure 5D). These pathways are hallmarks of Myc-driven transcriptional networks and emphasize the biological impact of this regulatory shift ^46,47^.

Strikingly, annotation of the Miz1-loss/Myc-gain associated target genes revealed that the corresponding regulatory regions were predominantly promoter-proximal, with ∼90% located within 1 kb of transcriptional start sites (Figure 5E). Finally, motif enrichment analysis using HOMER revealed that the Miz1-lost and Myc-gained regions are enriched for canonical E-box motifs (CACGTG), the preferred Myc binding sites (Figure 5F). Together, these findings suggest a model where loss of repressive Miz1 binding in RPMM tumors and moreover Myc recruitment to promoter regions results in activation of Myc-associated transcriptional programs (see Figure 1D, Supplementary Figure 1E, F). Furthermore, our data indicate that transcriptional activation in RPMM tumors is not driven by a global increase in Myc occupancy, but rather by a redistribution of Myc binding toward genes that also lose Miz1-associated regulatory input. At these shared target genes, loss of promoter-associated Miz1 coincides with retained or increased Myc occupancy, thereby relieving repression and favoring activation of Myc-associated transcriptional programs.

Prompted by the observation that loss of Miz1 and gain of Myc binding in RPMM tumors led to activation of RNA processing and apoptosis related programs - suggestive of increased cellular and mitochondrial stress - we next asked whether these transcriptional changes translate into heightened apoptotic sensitivity. To test this, we performed BH3 profiling in RPM and RPMM tumor-derived cells to functionally assess mitochondrial apoptotic priming. Strikingly, RPMM cells exhibited significantly higher cytochrome c release across nearly all peptides and BH3 inhibitors, consistent with increased apoptotic priming compared to RPM cells (p < 0.0001), indicating a global increase in apoptotic sensitivity (Figure 5G).

These findings suggest that Myc-driven transcriptional reprogramming in RPMM tumors, characterized by elevated RNA and protein biosynthesis, imposes a heightened stress burden on the mitochondrial network, thereby lowering the threshold for apoptosis. Thus, chromatin-level changes induced by the Miz1^ΔPOZ^ mutation not only reshape transcriptional programs but also render RPMM cells functionally more susceptible to apoptotic signals.

Given the central role of Myc in modulating cellular stress responses and apoptosis, we next explored the translational relevance of this phenotype by assessing sensitivity to standard-of-care chemotherapeutic agents used in SCLC. RPM and RPMM tumor-derived cell lines were treated with cisplatin and etoposide, the most widely used first-line chemotherapeutic for SCLC patients. While no significant difference in cisplatin sensitivity was observed between genotypes (p = 0.1294), RPMM cells exhibited significantly lower GI₅₀ values for etoposide compared to RPM cells (p = 0.0253), indicating increased sensitivity to etoposide specifically (Figure 5H).

To directly test whether the increased sensitivity to etoposide is driven by enhanced Myc activity following loss of DNA-bound Miz1, we employed an inducible Myc-ER construct to acutely elevate nuclear Myc levels in Miz1 unperturbed SCLC cell lines (Supplementary Figure 5A). Activation of Myc by 4-hydroxytamoxifen (OHT) markedly increased sensitivity to etoposide treatment, both in cells expressing wild-type Myc-ER and in those expressing the phosphorylation-deficient Myc^T58A^-ER mutant (Figure 5I). Consistent with this enhanced drug sensitivity, immunofluorescence staining of nuclear γH2AX, a marker of DNA damage, was significantly increased upon Myc^T58A^ nuclear translocation in etoposide-treated cells (Figure 5J). These findings support a model in which elevated Myc activity sensitizes tumor cells to etoposide-induced DNA damage, linking Myc-driven transcriptional states to chemotherapy responsiveness.

To characterize transcriptional responses to etoposide, we performed RNA-seq after 24 h and 72 h of treatment in RPM and RPMM cells. Etoposide induced a robust stress response in both genotypes, marked by downregulation of cell cycle-associated genes and upregulation of interferon and inflammatory signaling pathways, with effects becoming more pronounced at 72 h (Figure 5K, Supplementary Table 1). Gene set enrichment analysis further revealed repression of oxidative phosphorylation and Myc target gene signatures alongside activation of interferon-α and interferon-γ pathways (Figure 5K). To directly compare the transcriptional response between RPM and RPMM cells, we examined log2 fold-changes induced by etoposide in both genotypes. Remarkably, gene expression changes were highly correlated between RPM and RPMM cells at both 24 h and 72 h (Supplementary Figure 5B-C), indicating that etoposide elicits largely similar transcriptional programs irrespective of Miz1 status. These findings indicate that enhanced etoposide sensitivity in RPMM cells is not due to a distinct drug-induced transcriptional response, but rather reflects their pre-existing transcriptional rewiring and elevated apoptotic priming.

To evaluate the impact of the Miz1^ΔPOZ^ loss on treatment response in vivo, RPM and RPMM mice were treated with a standard-of-care combination of cisplatin and etoposide (Figure 5L). Kaplan-Meier survival analysis confirmed a significant overall benefit from chemotherapy, and a non-significant trend toward improved survival in RPMM mice compared to RPM was observed, suggesting a possible genotype-dependent difference in treatment response (Figure 5M). In line, serial MRI imaging of representative mice suggested a pronounced tumor shrinkage in RPMM mice treated with chemotherapy compared to RPM counterparts (Figure 5N). Consistent with this observation, tumor burden trajectories over time suggest a stronger reduction in chemotherapy-treated RPMM mice compared to RPM mice (Figure 5O). To quantitatively assess therapeutic response at the study endpoint, tumor size changes from baseline were summarized in a waterfall plot (Figure 5P). Statistical testing was performed using a bootstrap-based difference-in-differences approach (10,000 resamples), comparing the chemotherapy effect (Chemo–Vehicle) between RPM and RPMM genotypes. This analysis demonstrated a significantly stronger treatment effect in RPMM tumors (DOD = −1282; 95% bootstrap confidence interval −2782 to −20; p = 0.046). Further quantifying the therapeutic response, mice were classified using RECIST-like criteria into progressive disease (PD), stable disease (SD), partial response (PR), and complete response (CR). This analysis revealed a stark contrast: all RPM mice were either PD (progressive disease) or PR (partial response; eight and two mice, respectively), with no cases of SD (stable disease) or CR (complete response). In contrast, RPMM tumors showed superior response, with three PR and two CR (Figure 5P). These data underscore the nuanced effects of Myc protein distribution at distinct chromosomal sites and the direct impact of Myc activity levels on genotoxic stress and chemosensitivity in SCLC tumors.

## Discussion

SCLC is defined by an aggressive neuroendocrine lineage state, yet tumors frequently undergo transcriptional rewiring and dedifferentiation during progression and therapy resistance ^48,49^. The transcription factor MYC has been implicated in these state transitions ^8,13,50^, but how its chromatin activity is regulated in lineage-specific contexts remains incompletely understood. Here, we identify the zinc-finger protein Miz1 as a critical determinant of MYC’s genomic binding pattern in SCLC. Using a novel Miz1^ΔPOZ^-driven RPMM mouse model, we demonstrate that loss of Miz1 DNA binding reshapes Myc chromatin occupancy between enhancers and promoters, thereby coordinating two opposing transcriptional outcomes: silencing of neuroendocrine differentiation programs and activation of MYC-driven stress and apoptotic pathways. In parallel, integrative analyses in MYC-amplified human SCLC cell lines and patient-derived specimens show that this enhancer architecture, including low-affinity E-boxes, is conserved and preferentially associated with neuroendocrine programs, underscoring the relevance of this mechanism in human disease. Collectively, these findings establish Miz1 as a context-dependent chromatin regulator that constrains Myc transcriptional output at promoters while simultaneously enabling Myc binding at lineage-specific enhancers in both murine and human SCLC.

Miz1 has classically been linked to promoter-associated repression within Myc networks, particularly at genes controlling differentiation and cell cycle arrest ^16,19,51,52^. Our findings broaden this view by revealing that Miz1 also plays an essential role at distal regulatory elements, where it supports Myc occupancy at enhancers governing neuroendocrine lineage programs. This context-dependent function suggests that Miz1 shapes Myc activity by directing its genomic association, as loss of Miz1 DNA occupancy leads to diminished Myc binding at lineage-maintaining enhancer circuits and a concomitant redistribution toward promoter-associated regions, thereby enhancing stress and biosynthetic transcriptional programs. Mechanistically, regions with cooperative loss of Miz1 and Myc binding show reduced chromatin accessibility, decreased H2A.Z and H3K36me3 deposition, and diminished eRNA production, together indicating that Miz1/Myc-bound enhancers constitute genuinely active regulatory elements whose function is silenced upon Miz1^ΔPOZ^ mutation. In contrast, promoter regions with loss of Miz1 and gain or maintenance of Myc occupancy are linked to genes involved in ribosome biogenesis, RNA processing, and apoptosis, suggesting that removal of promoter-bound Miz1 allows canonical Myc target programs to become more fully engaged.

Such enhancer-directed cooperativity is consistent with emerging models in which the oncogenic output of Myc is critically dependent on regulatory architecture beyond promoters ^15,43^. While Miz1 has long been studied as a transcriptional regulator ^53–55^, enhancer-associated Miz1 DNA binding has remained largely unexplored. Our data suggest that Miz1 can stabilize Myc recruitment at suboptimal enhancer sites. Notably, low-affinity E-boxes are enriched at distal regulatory regions in both mouse tumors and human SCLC models, whereas canonical E-boxes remain promoter biased, implying that low-affinity motifs may serve as a general mechanism to encode lineage specificity into Myc-dependent enhancer usage. Consistent with this, we observe a distributed set of Miz1/Myc-bound enhancers rather than a discrete super-enhancer class, arguing that a network of cooperative enhancer elements, rather than a small number of focal hubs, underlies maintenance of the neuroendocrine state in MYC-driven SCLC. This motif-based mechanism offers a framework for how lineage-specific enhancer architecture shapes Myc-dependent transcriptional programs in SCLC.

Our data are compatible with a model in which, in a baseline state with low neuroendocrine signaling, Myc predominantly engages high-affinity E-boxes at promoters to sustain biosynthetic and cell cycle programs, whereas low-affinity E-boxes at distal neuroendocrine enhancers remain largely inactive. Upon Myc amplification, excess Myc can be recruited to these suboptimal E-boxes, where Miz1 binding stabilizes Myc occupancy at enhancers located near lineage-defining transcription factors such as ASCL1 and NEUROD1, as well as their downstream neuroendocrine targets. This suggests a cooperative mechanism in which Myc, Miz1, and core neuroendocrine factors jointly establish and maintain neuroendocrine enhancer circuits, potentially explaining why the genomic regions that lose both Miz1 and Myc in RPMM tumors are enriched for motifs of neuroendocrine transcription factors. At the same time, the mutual exclusivity of MYC amplification and ASCL1-high SCLC ^56^, and the relatively low neuroendocrine signaling in MYC-amplified subtypes, indicate that Myc may also function as a secondary or compensatory regulator that maintains a baseline neuroendocrine gene expression state when core lineage factors such as ASCL1 are diminished, rather than fully substituting for them. Consistent with this idea, MYC family amplification is frequently observed across neuroendocrine malignancies, including SCLC ^8^, neuroendocrine prostate cancer ^57^, medulloblastoma ^58^, and pancreatic ductal adenocarcinoma ^59^, raising the possibility that context-dependent engagement of low-affinity E-boxes at lineage enhancers represents a recurrent mechanism of Myc-mediated neuroendocrine plasticity.

An important implication of our findings is that Miz1-dependent control of Myc chromatin binding may represent a broader regulatory principle beyond SCLC. Consistent with this, our analysis of an independent Miz1^ΔPOZ^ lymphoma model ^20^ revealed similarly increased activation of Myc target signatures, suggesting that enhanced Myc-driven transcription upon loss of Miz1 DNA binding is conserved across tumor entities. Strikingly, however, the suppressed transcriptional programs were highly lineage-specific, affecting B-cell-associated pathways rather than neuroendocrine differentiation ^60^. These observations support a model in which Miz1 enables Myc occupancy at cell type-restricted enhancer circuits that maintain lineage-defining transcriptional states. Together with our SCLC data, this suggests that Miz1 functions as a lineage-sensitive gatekeeper that couples a broadly acting oncogenic factor to cell type-specific enhancer logic. Future studies will be required to determine how broadly enhancer-directed Miz1-Myc cooperativity operates across malignancies and whether it can be therapeutically exploited.

A particularly unexpected implication of this model is that Myc activation and lineage loss can coexist with increased apoptotic priming and chemotherapy sensitivity. Myc is well known to impose intrinsic cellular stress, creating a state of heightened dependence on mitochondrial survival pathways ^9,61–63^. Our findings suggest that when Miz1-mediated repression at promoters is relieved, enabling a concomitant gain of Myc occupancy, Myc-driven biosynthetic and stress programs become amplified, lowering the apoptotic threshold and sensitizing tumors to DNA damaging agents such as etoposide. In parallel, cooperative loss of Miz1/Myc binding at lineage-maintaining enhancers reduces neuroendocrine gene expression and cellular heterogeneity, indicating that the same tumor cells can simultaneously undergo lineage remodeling and experience heightened metabolic and replicative stress. This may help reconcile the paradox that MYC-high SCLC can display both aggressive dedifferentiation and enhanced treatment vulnerability, consistent with recent studies linking transcriptional subtype to therapeutic response ^9,13,64,65^.

From a translational perspective, our findings suggest that Myc-driven lineage plasticity in SCLC may create specific therapeutic vulnerabilities rather than uniformly promoting treatment resistance. Loss of Miz1 DNA binding shifts Myc activity toward promoter-associated stress and biosynthetic programs, resulting in increased mitochondrial apoptotic priming and enhanced sensitivity to etoposide. These data raise the possibility that chromatin states defined by Miz1-Myc enhancer versus promoter occupancy could influence chemotherapy responsiveness in MYC-driven SCLC. More broadly, therapeutic strategies that exploit Myc-induced apoptotic stress, including DNA-damaging agents or apoptosis-sensitizing approaches, may be particularly effective in tumors that lose neuroendocrine lineage differentiation.

Future work will be needed to dissect how Miz1 promotes Myc binding at lineage-specific enhancers. It will also be important to define which additional cofactors, including neuroendocrine lineage transcription factors such as ASCL1 or NEUROD1 suggested by motif enrichment, physically contribute to Miz1/Myc complexes at enhancers. Beyond transcription factor co-occupancy, identifying the chromatin-modifying complexes whose activity maintains H2A-Z deposition and H3K36me3 at Miz1/Myc-bound enhancers will be essential to fully resolve the downstream effector mechanism. Ultimately, establishing the relevance of this regulatory logic in human SCLC and patient-derived settings will be critical. Moreover, extending our observations of increased apoptotic priming and etoposide sensitivity to additional therapeutic contexts may help define how Myc-driven stress states can be exploited for improved treatment strategies.

## Acknowledgements

We would like to thank all current and previous Sos Lab members, especially Marcel Dammert and Shaliny Sothyratnam as well as Rasmus Siersbaek, Alexander Sasse, and Martin Eilers for fruitful discussions. Also, we would like to thank Roman K. Thomas, Christian Reinhardt, Graziella Bosco and all members of the consortium for the highly cooperative environment in the SFB1399.

## Author contributions: CRediT

- Conceptualization: M.L.S
- Data curation: L.M.F., H.L.T., B.A., M.R., J.O., P.Z., P.H., M.H.W., Y.T., D.P.,
- Formal analysis: L.M.F., H.L.T., B.A., P.Z., P.H., M.H.W., D.P., J.F, F.B., J.B.
- Funding acquisition: M.L.S.
- Investigation: L.M.F., H.L.T., B.A., M.R., L.W., M.F., A.H., J.O., P.Z., P.H., M.H.W., Y.T, D.P.,
- Methodology: L.M.F., H.L.T., B.A., P.Z., P.H., M.H.W., Y.T., J.H., O.W., D.P., J.F, E.W.
- Project administration: K.G.
- Resources: E.W.
- Supervision: J.F., R.H.H, M.L.S.
- Visualization: L.M.F., P.Z.,
- Writing – original draft: L.M.F., M.L.S
- Writing – review and editing: L.M.F., M.L.S

## Supplemental Information

Document Supplementary_Figures. Figures S1–S5.

Table S1. GSEA.

Table S2. GO term enrichment.

Table S3. ABC.

Table S4. ROSE.

Table S5. AlphaGenome.

