## Supplementary_Figures for "MYC-MIZ1 Complexes at Enhancers Tune Neuroendocrine Identity of Small Cell Lung Cancer"

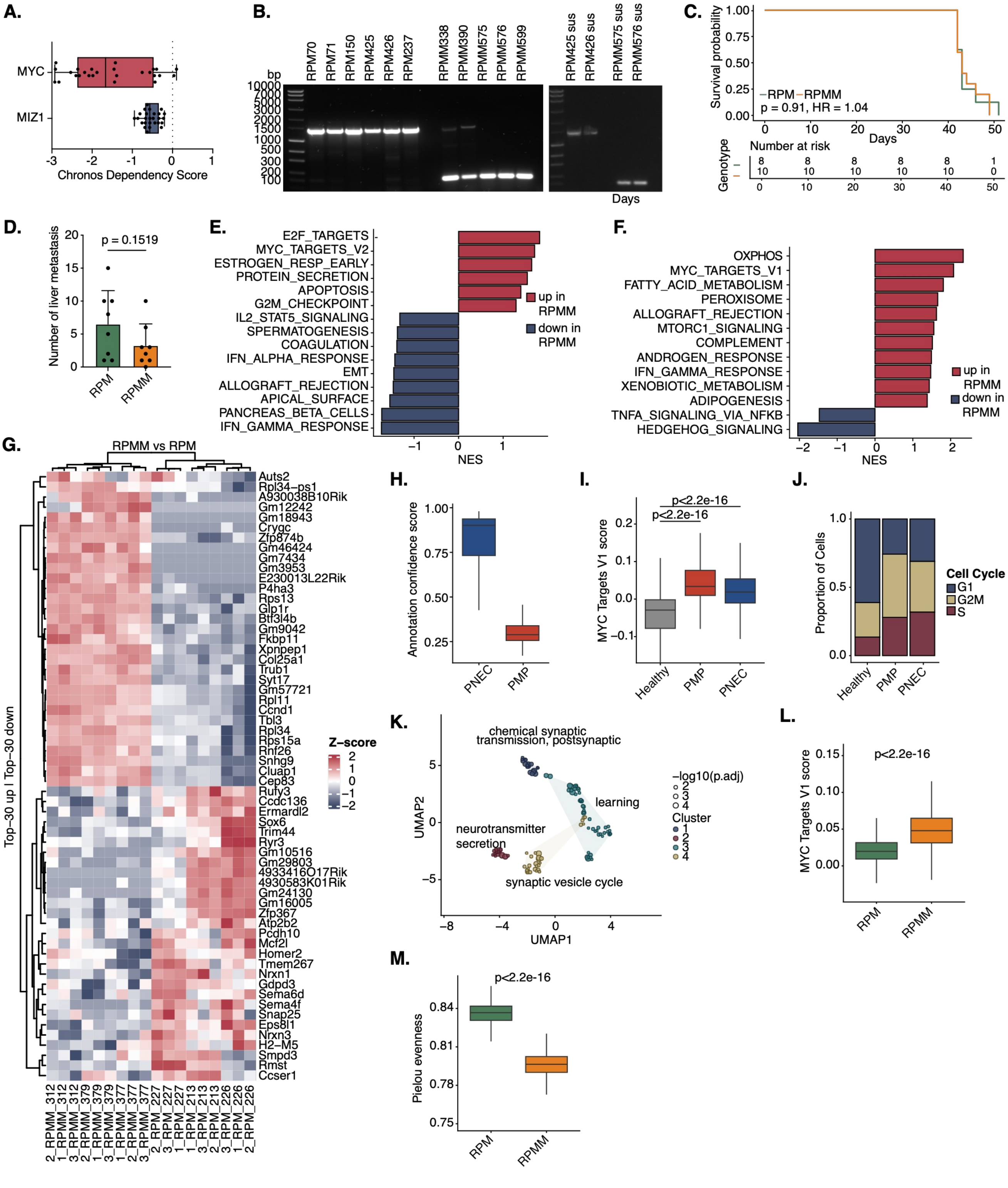


**Supplementary Figure 1: Miz1ΔPOZ Mutation Alters Tumor Characteristics of RPM Mouse Model**

**A.** Distribution of MYC and MIZ1 chronos dependency scores lung neuroendocrine tumors obtained from the DepMap portal data. Each point represents an individual human cell line. **B.** PCR-based genotyping of RPM and RPMM cell lines to assess Miz1 status. PCR was performed using primer 1 and 5, yielding an expected amplicon of ~1450 bp in RPM cell lines and ~200 bp amplicon in RPMM cell lines, consistent with deletion of the Miz1 POZ domain. **C.** Kaplan Meier Analysis of mice bearing RPM and RPMM tumors. Time after infection with CMV-Cre virus is shown. Survival differences between RPM and RPMM tumors were evaluated using the Mantel-Cox log-rank test. **D.** Number of liver metastases in RPM and RPMM mice is shown. T-test was used to compare RPM with RPMM mice. **E.-F.** GSEA of Hallmark pathways of 3'UTR RNA-seq of RPM (6 biological and 2 technical replicates) and RPMM cell lines (5 biological and 2 technical replicates; E.) and snRNA-seq of one RPM and two RPMM tumors (F.). Pathways with p-value < 0.05 were included. **G.** Top 30 up- and downregulated genes of 3'UTR bulk RNA sequencing of FFPE tissue isolated from RPM and RPMM tumors is shown. Gene expression is scaled per gene (Z-score). **H.** Cell-type specific annotation confidence score of RPM and RPMM snRNA-seq data in PNEC vs PMP cell types. **I.** Boxplots displaying the distribution of Hallmark MYC_TARGETS_V1 scores for snRNA-seq data from RPM and RPMM tumors, distributed in healthy (immune and endothelial cells), PMP and PNEC cells. Statistical significance was assessed using a Wilcoxon rank-sum test. **J.** Cell cycle phase distribution of tumor-associated populations (PNEC and PMP) compared with the non-malignant reference (immune + endothelial cells), shown as proportions of cells per group. **K.** Differential gene expression between PMP and PNEC tumor cell populations in snRNA-seq. Pseudobulk DESeq2 analysis comparing PMP versus PNEC cells and filtered for all significantly downregulated genes (padj < 0.05, |log2FC| ≥ 1) in PMP tumor cell populations. Significantly enriched GO terms were clustered by semantic similarity and shown in a UMAP plot. **L.** Boxplots displaying the distribution of Hallmark MYC_TARGETS_V1 scores for spatial transcriptomics data from RPM and RPMM tumors. Statistical significance was assessed using a Wilcoxon rank-sum test. **M.** Boxplot of Pielou's evenness analysis of pseudotime distributions. Pseudotime was inferred as described in Figure 1 N, scaled to 0-1 within each genotype, and divided into equal-width bins. Pielou evenness was calculated from bin occupancy. Statistical significance was assessed using a Wilcoxon rank-sum test on bootstrapped data.

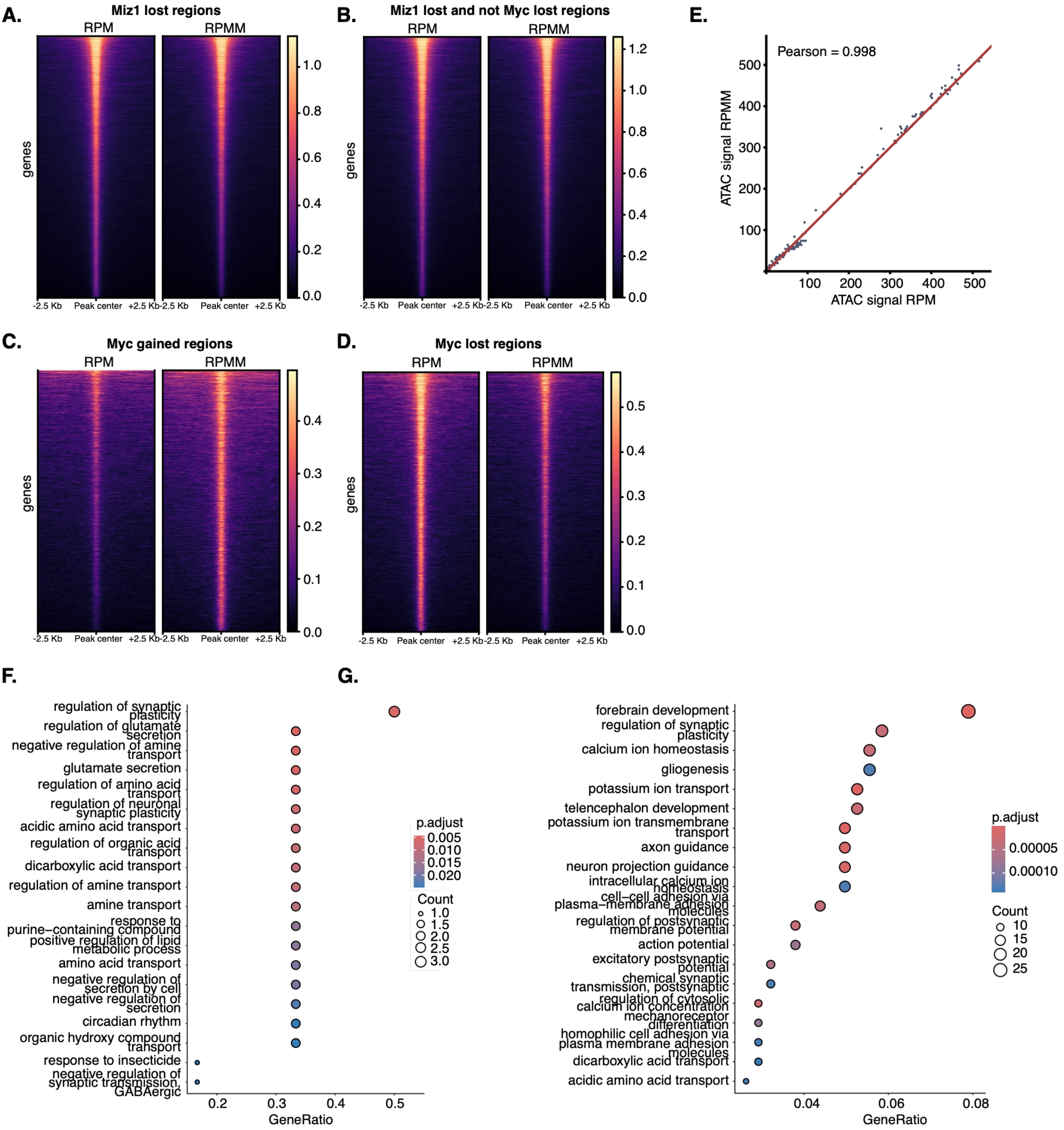


**Supplementary Figure 2: Cooperative Loss of Miz1 and Myc on Chromatin Defines a Distinct Transcriptional Subprogram**
**A.-D.** Heatmaps of ATAC signal intensity in RPM and RPMM tumors corresponding to the average profiles shown in Figure 2D. **E.** Genome-wide comparison of ATAC-seq signal between RPM and RPMM tumors. ATAC-seq signal intensities were summarized in 100-bp genomic bins and plotted as a hexbin scatter plot. **F.-G.** GO term enrichment of all significantly downregulated genes ($log_2FC<-1$, FDR<0.05) in the RPMM model. 3'UTR bulk RNA-seq data of RPM (6 biological and 2 technical replicates) and RPMM (5 biological and 2 technical replicates) cell lines (F.) or snRNA-seq of one RPM and two RPMM tumors (G.) was used. Top 20 enriched GO terms are shown.

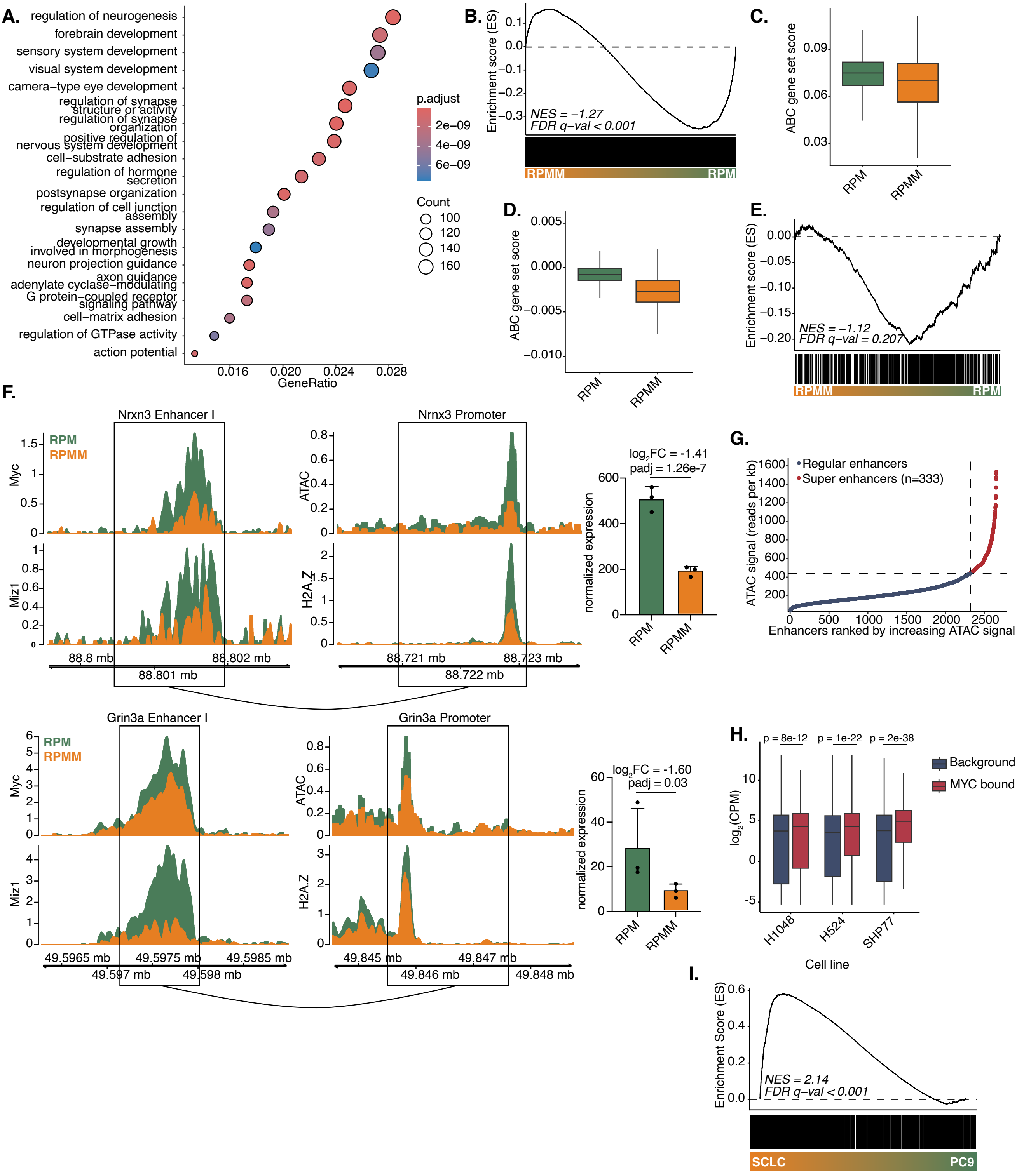


**Supplementary Figure 3: Cooperative Binding of Miz1 and Myc at Enhancers Maintains Neuroendocrine Differentiation**

**A.** GO term enrichment of the 6,790 unique target genes of the ABC output using 3'UTR bulk RNA-seq of FFPE tissue isolated from RPMM and RPM tumors. **B.** GSEA of 6,790 unique target genes of the ABC model using 3'UTR bulk RNA-seq of RPM and RPMM cell lines. **C.-D.** Module score of the 6,790 unique target genes of the ABC model using single cell RNA-seq (C.) and spatial transcriptomics (D.) data of RPM and RPMM tumors. Statistical significance was assessed using a Wilcoxon rank-sum test. **E.** GSEA of 1,479 genes highly expressed in SCLC and linked to neuroendocrine signaling using 3'UTR bulk RNA-seq of FFPE tissue isolated from RPMM and RPM tumors. **F.** Signal intensity of Miz1 and Myc (left) as well as ATAC and H2A.Z (right) in RPM (green) and RPMM (orange) tumors. The ABC-predicted enhancer and promoter regions of Nrxn1 (upper graphic) and Grin3a (lower graphic) are shown. The bar plot represents the normalized RNA expression of the respective gene using 3'UTR bulk RNA-seq of FFPE tissue isolated from RPMM and RPM tumors. $Log_2FC$ and padj were obtained from the DESeq2 differential expression analysis. **G.** ROSE-style ranking of ABC output regions by ATAC-seq signal in RPM tumors. ABC output regions were stitched (within 12.5 kb, excluding ±2 kb around transcription start sites) and ranked by chromatin accessibility. Super-enhancers (red) were defined based on the inflection point in ATAC-seq signal intensity, with regular enhancers shown in blue. Dashed lines indicate the signal threshold used for super-enhancer classification. **H.** Expression of genes linked to MYC-bound enhancers compared with the ABC-linked background in human MYC-amplified SCLC cell lines. Analysis was restricted to 6,790 unique target genes derived from ABC enhancer regions corresponding to the murine Myc and Miz1 lost regions in RPMM tumors. Gene expression ($\log_2$(CPM)) is shown for genes assigned by the human ABC model to MYC-bound enhancers (red) and for all genes linked by the ABC model within this gene set, irrespective of MYC binding (background, blue). Statistical significance was assessed using a wilcoxon rank-sum test. **I.** GSEA of the 1,479 genes highly expressed in SCLC and linked to neuroendocrine signaling from the murine model, mapped to human orthologs, using 3'UTR bulk RNA-seq from human SCLC cell lines (SCLC21H and DMS273) compared to the human NSCLC cell line PC9.

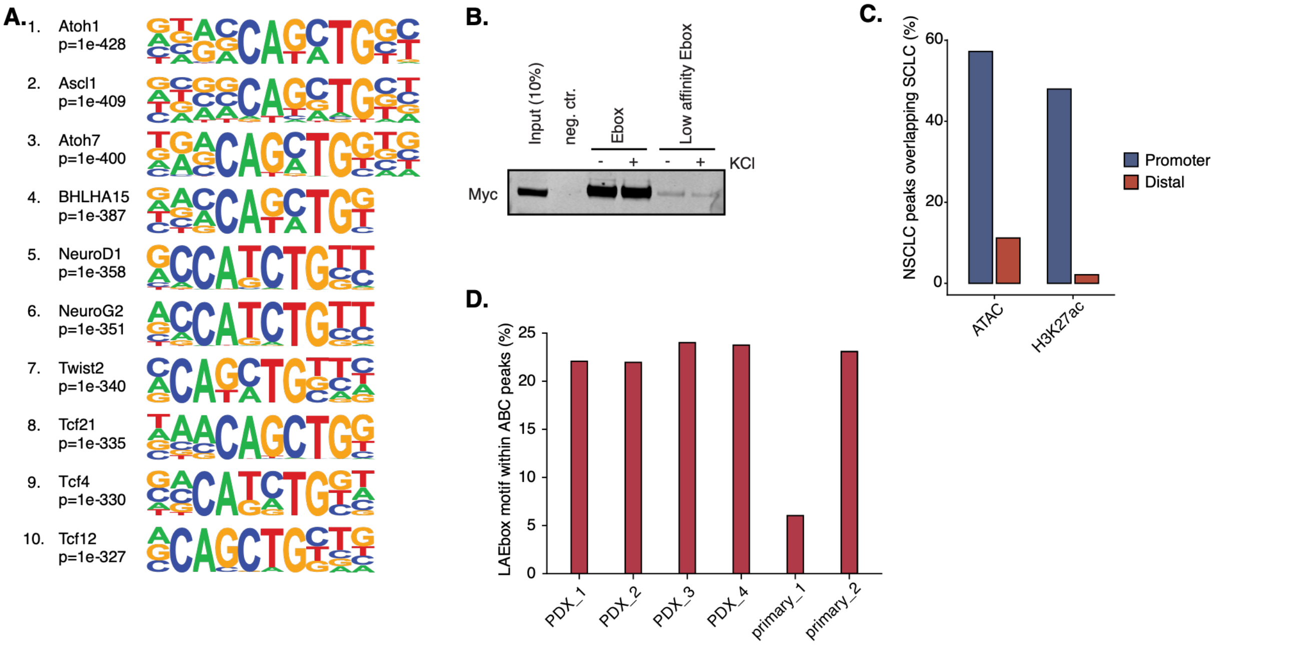


**Supplementary Figure 4: Cooperative Miz1–Myc DNA binding at enhancers is mediated by a novel Miz1 DNA binding motif**

**A.** HOMER motif enrichment analysis of Miz1 and Myc lost regions in RPMM tumors, showing top ten enriched motifs. **F.** Western blot analysis of DNA pulldown assays. Myc binding to the canonical E-box (CACGTG) and the low-affinity E-box (CAGCTG) using RPM150 cells. Washing was performed with 110 mM KCl (-) and 300 mM KCl (+). **C.** Percentage of NSCLC peaks overlapping SCLC consensus peaks across ATAC-seq and H3K27ac DynaTag. **D.** Enrichment of the low-affinity E-box (LAEbox) motif within ABC-defined enhancer regions in human primary FFPE biopsies (primary) and PDX samples (PDX) from SCLC patients. Bar plot showing the fraction of ABC regions containing the LAEbox motif.

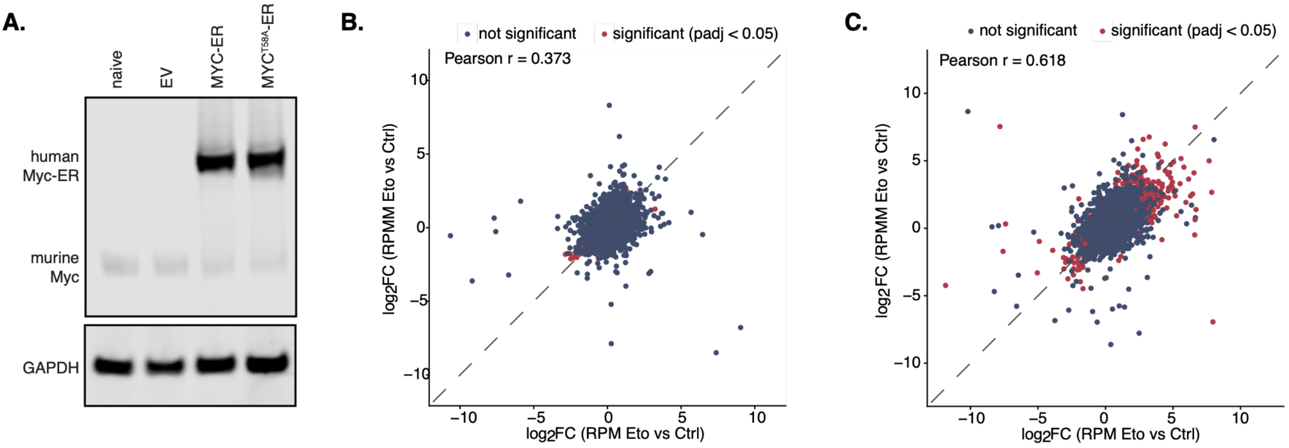


**Supplementary Figure 5: Promoter-Localized Loss of Miz1 Relieves Repression of Myc-Driven Transcriptional Programs and Increases Chemotherapeutic Sensitivity**
**A.** Western Blot of RP1380 cells stably transduced with empty vector (EV), MYC-ER, or MYC$^{T58A}$-ER constructs. **B.-C.** Scatter plot of $log_2FC$ (etoposide vs. control) derived from 3'UTR RNA-seq for RPM (x-axis) and RPMM (y-axis) cell lines after 24h (A.) or 72h (B.) of treatment. Each dot represents one gene. Genes significantly differentially expressed in at least one genotype (DESeq2, padj < 0.05) are shown in red, non-significant genes in blue. The dashed line indicates the line of identity (y=x). Pearson correlation coefficients (r) are indicated in each panel.
